# How motivational signals disrupt metacognitive signals in the human VMPFC

**DOI:** 10.1101/2020.10.02.323550

**Authors:** Monja Hoven, Gina Brunner, Nina de Boer, Anneke Goudriaan, Damiaan Denys, Ruth van Holst, Judy Luigjes, Mael Lebreton

**Affiliations:** Department of Psychiatry, Amsterdam UMC, University of Amsterdam, Amsterdam, The Netherlands; Arkin and Jellinek, Mental Health Care, Amsterdam, The Netherlands; Netherlands Institute for Neuroscience, an Institute of the Royal Netherlands Academy of Arts and Sciences, Amsterdam, The Netherlands; Swiss Center for Affective Science, University of Geneva, Switzerland; Laboratory for Behavioral Neurology and Imaging of Cognition, Department of Fundamental Neurosciences, University of Geneva, Switzerland

**Keywords:** decision-making, VMPFC, confidence, incentives, motivation, bias, fMRI

## Abstract

A growing body of evidence suggests that, during decision-making, BOLD signal in the VMPFC correlates both with motivational variables – such as incentives and expected values – and metacognitive variables – such as confidence judgments, which reflect the subjective probability of being correct. At the behavioral level, we recently demonstrated that the value of monetary stakes bias confidence judgments, with gain (respectively loss) prospects increasing (respectively decreasing) confidence judgments, even for similar levels of difficulty and performance. If and how this value-confidence interaction is also reflected in VMPFC signals remains unknown. Here, we used an incentivized perceptual decision-making task that dissociates key decision-making variables, thereby allowing to test several hypotheses about the role of the VMPFC in the incentive-confidence interaction. While initial analyses seemingly indicate that VMPFC combines incentives and confidence to form an expected value signal, we falsified this conclusion with a meticulous dissection of qualitative activation patterns. Rather, our results show that strong VMPFC confidence signals observed in trials with gain prospects are disrupted in trials with no – or negative (loss) monetary prospects. Deciphering how decision variables are represented and interact at finer scales (population codes, individual neurons) seems necessary to better understand biased (meta)cognition.

## Introduction

Over the past decades, a growing number of neurophysiological studies in human and non-human primates have established that the neural signals recorded during learning and decision-making tasks in the orbito-medial parts of the prefrontal cortex (OMPFC) – the medial orbitofrontal cortex (OFC) and the ventromedial prefrontal cortex (VMPFC) – correlate with key variables derived from theories of motivation and decision-making (Kable and Glimcher, 2009; Padoa-Schioppa, 2007; Rangel and Hare, 2010). For instance, in Pavlovian conditioning tasks, neurons in the non-human primate OFC signal the anticipatory value of upcoming rewards, with neural activity commensurate to the monkeys’ subjective preferences (Tremblay and Schultz, 1999). In economic decision-making tasks, neuronal activity in the same region of the OFC correlates with the subjective value of available options (Padoa-Schioppa and Assad, 2006). In humans, similar results have been derived from functional neuroimaging studies. Blood-Oxygen Level Dependent (BOLD) signal in the VMPFC scales with the anticipation of upcoming reward (Kahnt et al., 2011; Knutson et al., 2003), with the subjective pleasantness and desirability attributed to different stimuli (Lebreton et al., 2009), with the willingness to pay for different types of goods (Chib et al., 2009; Levy and Glimcher, 2011; Plassmann et al., 2007), and with the expected value of prizes, performance incentives and economic bundles such as lotteries (Gläscher et al., 2009; Hare et al., 2008; Knutson et al., 2005; McNamee et al., 2013). Overall, together with the midbrain and the ventral striatum (VS), the VMPFC seems to form a Brain Valuation System (Bartra et al., 2013; Haber and Behrens, 2014; Haber and Knutson, 2009), whose activity automatically indexes the value of available options so as to guide value-based decision-making (Lebreton et al., 2009; Levy et al., 2011), and to motivate motor and cognitive performance (Pessiglione and Lebreton, 2015).

Recently, a set of human neurophysiological studies have suggested that activity in the VMPFC is also triggered by metacognitive processes (Fleming and Dolan, 2012; Vaccaro and Fleming, 2018). In particular, both single neurons and BOLD activity in the VMPFC correlate with participants’ confidence in their own judgments and choices (De Martino et al., 2013; Lebreton et al., 2015; Lopez-Persem et al., 2020; Shapiro and Grafton, 2020). Confidence is a metacognitive variable that can be defined as one’s subjective estimate of the probability of being correct (Fleming and Daw, 2017; Pouget et al., 2016). Just like values, confidence judgments seem to be automatically represented in the VMPFC, for different types of judgments and choices (Lebreton et al., 2015; Lopez-Persem et al., 2020; Morales et al., 2018). Confidence signals are potentially useful to the flexible adjustment of behavior – such as monitoring and reevaluating previous decisions (Folke et al., 2016), tracking changes in the environment (Heilbron and Meyniel, 2019; Vinckier et al., 2016), or arbitrating between different strategies (Daw et al., 2005; Donoso et al., 2014).

Interestingly, at the behavioral level, values and confidence seem to interact. For instance, a handful of studies in psychology and economics have documented that confidence judgments increase in response to prospects of monetary bonuses (Giardini et al., 2008), and to positive incidental psychological states such as elevated mood (Koellinger and Treffers, 2015), absence of worry (Massoni, 2014) and emotional arousal (Allen et al., 2016; Jönsson et al., 2005). Recently, we designed an incentivized perceptual decision-making task to demonstrate that monetary incentives bias confidence judgments, with gain (respectively loss) prospects increasing (respectively decreasing) confidence judgments, even for similar levels of difficulty and performance (Lebreton et al., 2018). This result was also replicated in a reinforcement-learning context (Lebreton et al., 2019a; Ting et al., 2020). We explicitly hypothesized that this interaction would stem from the concurrent neural representation of – hence putative interaction between – incentives and confidence in the VMPFC (Lebreton et al., 2018).

Here, we use a functional neuroimaging adaptation of our original perceptual decision-making paradigm that allows to investigate the overlap in neural correlates between incentive and confidence (Lebreton et al., 2018). Our first set of analyses did not show the hypothesized overlap of incentive and confidence signals in the VMPFC at the expected statistical threshold (P < 0.05 whole-brain corrected FWE at the cluster level), nor in other regions of interest (ROI) that have been linked with value, motivation and confidence in the past - such as the ventral striatum (VS) and the anterior cingulate cortex (ACC). Therefore, we formulated an alternative hypothesis, positing that VMPFC integrates confidence and incentive signals into an expected value signal. We ran several quantitative and qualitative analyses that thoroughly compare the relative merits of these different hypotheses for the neural basis of the incentive-confidence interaction. Our results ultimately depict a complex picture, suggesting that motivational signals (notably prospects of loss) can disrupt metacognitive signals in the VMPFC.

## Results

To investigate the neurobiological basis of the interactions between incentives and confidence, we modified the task used in (Lebreton et al., 2018) to make it suitable for fMRI (**Figure 1A**). Basically, this task is a simple perceptual task (contrast discrimination), featuring a 2-alternative forced choice followed by a confidence judgment. At each trial, participants can win (gain context) or lose (loss context) points – or not (neutral context) – depending on the correctness of their choice. Importantly, this incentivization was implemented after the moment of choice and before confidence rating. Note that this corresponds to the simplest implementation of the task – corresponding to Experiment 2 in (Lebreton et al., 2018) –, which otherwise conditioned monetary outcomes to confidence rating precision rather than choice accuracy – see (Lebreton et al., 2018) for details. Yet, our previous results suggested that this task still reveals an effect of incentives on confidence, while keeping instructions simpler – a most desirable feature especially for clinical and fMRI studies.

**Figure 1.**
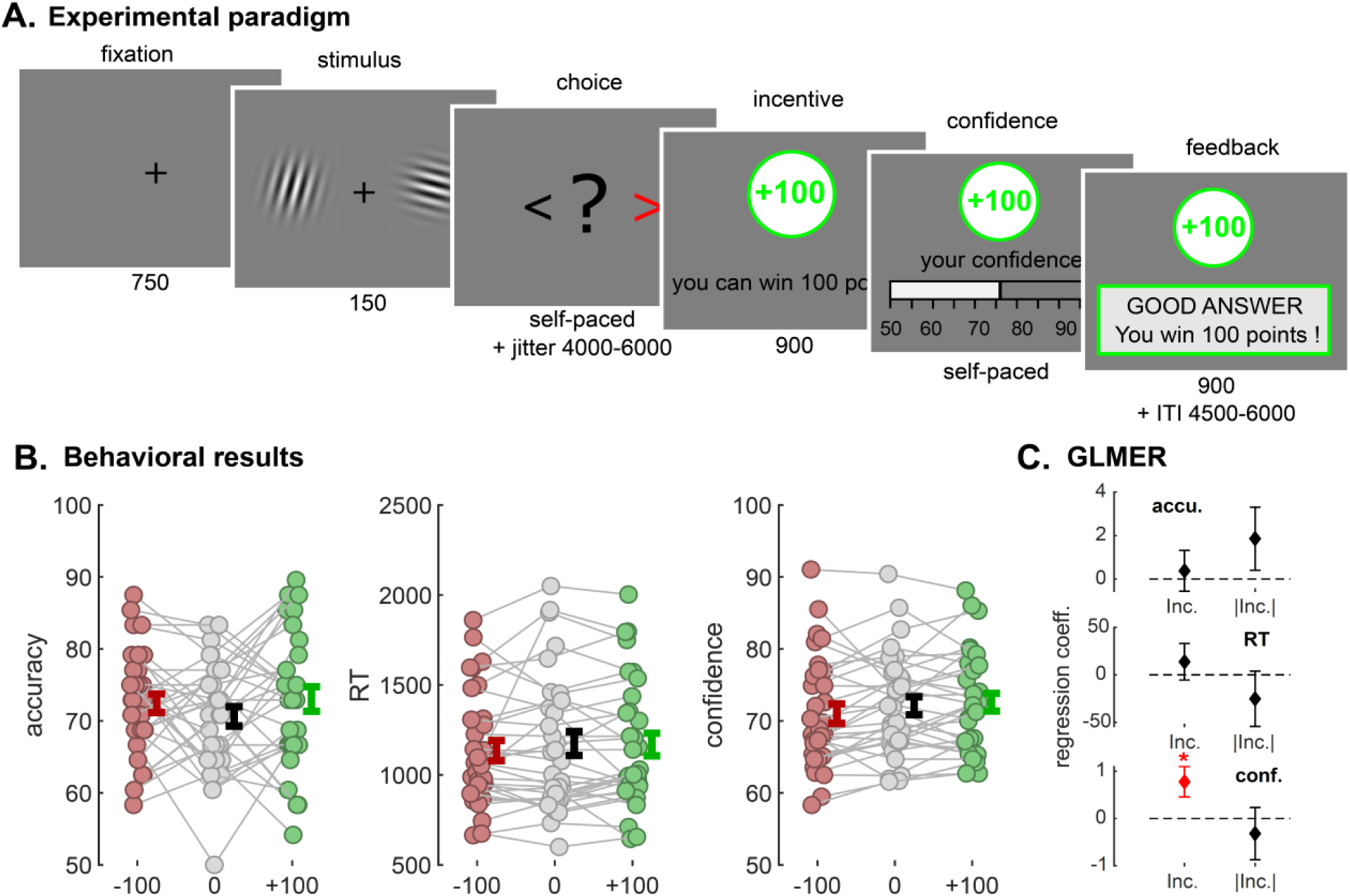
**A**. Experimental paradigm. Participants viewed two Gabor patches on both sides of the screen (150 ms) and then chose which had the highest contrast (left/right, self-paced). After a jitter of a random interval between 4500 to 6000 ms, the incentive was shown (900 ms; green frame for win trials, grey frame for neutral trials, red frame for loss trials). Then participants were asked to report their confidence in the earlier made choice on a scale ranging from 50% to 100% with steps op 5%. The initial position of the cursor was randomized between 65% and 85%. Finally, subjects received feedback. The inter trial interval (ITI) had a random duration between 4500 and 6000 ms. The calibration session only consisted of Gabor discrimination, without confidence rating, incentives or feedback and was used to adjust difficulty so that every individual reached a performance of 70%. **B**. Behavioral results. Individual-averaged accuracy (left), reaction times (middle) and confidence (right) as a function of incentive condition (−100/red, 0/grey, +100/green). Colored dots represent individuals, grey lines highlight within subject variation across conditions. Error bars represent sample mean ± sem. Note that for confidence and accuracy, we computed the average per incentive level per individual, but that for reaction times, we computed the median for each incentive condition rather than the mean due to their skewed distribution. **C**. Generalized linear mixed-effect regression (GLMER) results. Graph depict fixed-effect regression coefficients (β) for incentive net value and incentive absolute value predicting performance (top), reaction-times (middle) and confidence (bottom). Error bars represent standard errors of fixed effects. * P < 0.05

### Behavioral results replicate Lebreton, et al. 2018

To begin with, we verified that our task generated the incentive-confidence interaction at the behavioral level. First, using an approach similar to (Lebreton et al., 2018), we simply tested the biasing effects (i.e. net value) and motivational effects (i.e. absolute value) of incentives on behavioral variables, using linear mixed-effects models (see **Methods**). Replicating our previous results, we found a significant positive effect of incentive net value on confidence (β = 0.77 ± 0.32, t_4317_ = 2.43, P = 0.015) and no effect of incentive absolute value (β = −0.32 ± 0.55, t_4317_ = −0.58, P = 0.565; **Figure 1B**). This result alone validates the presence of incentive-confidence interaction at the behavioral level. Importantly, this effect was not driven by any net incentive value effects on accuracy or RT (accuracy: β = 0.38 ± 0.93, t_4317_ = 0.41, P = 0.685; RT: β = 13.75 ± 19.22, t_4317_ = 0.72, P = 0.474). Moreover, we did not find evidence for any absolute incentive value on neither accuracy nor RT (accuracy: β = 1.86 ± 1.45, t_4317_ = 1.28, P = 0.199; RT: β = −25.24 ± 29.17, t_4317_ = −0.87, P = 0.387). To confirm the robustness of our main effect of net incentive on confidence, we next ran several full linear mixed-effects models, which included additional control variables (evidence, accuracy, etc. see **Suppl. Mat**.). Overall, the incentive-confidence interaction remained significant after accounting for those other potential sources of biases/confounds.

### fMRI results

Having established the presence of a robust confidence-incentive interaction at the behavioral level, we next turned to the analysis of the functional neuroimaging data. Critically, our task allows us to temporally distinguish the moment of stimulus presentation and choice – where the decision value, together with an implicit estimation of (un)certainty are expected to build up – from the incentive presentation and confidence rating moment – where the explicit, metacognitive confidence signal is expected to interact with the incentive (**Figure 2A-B**).

**Figure 2.**
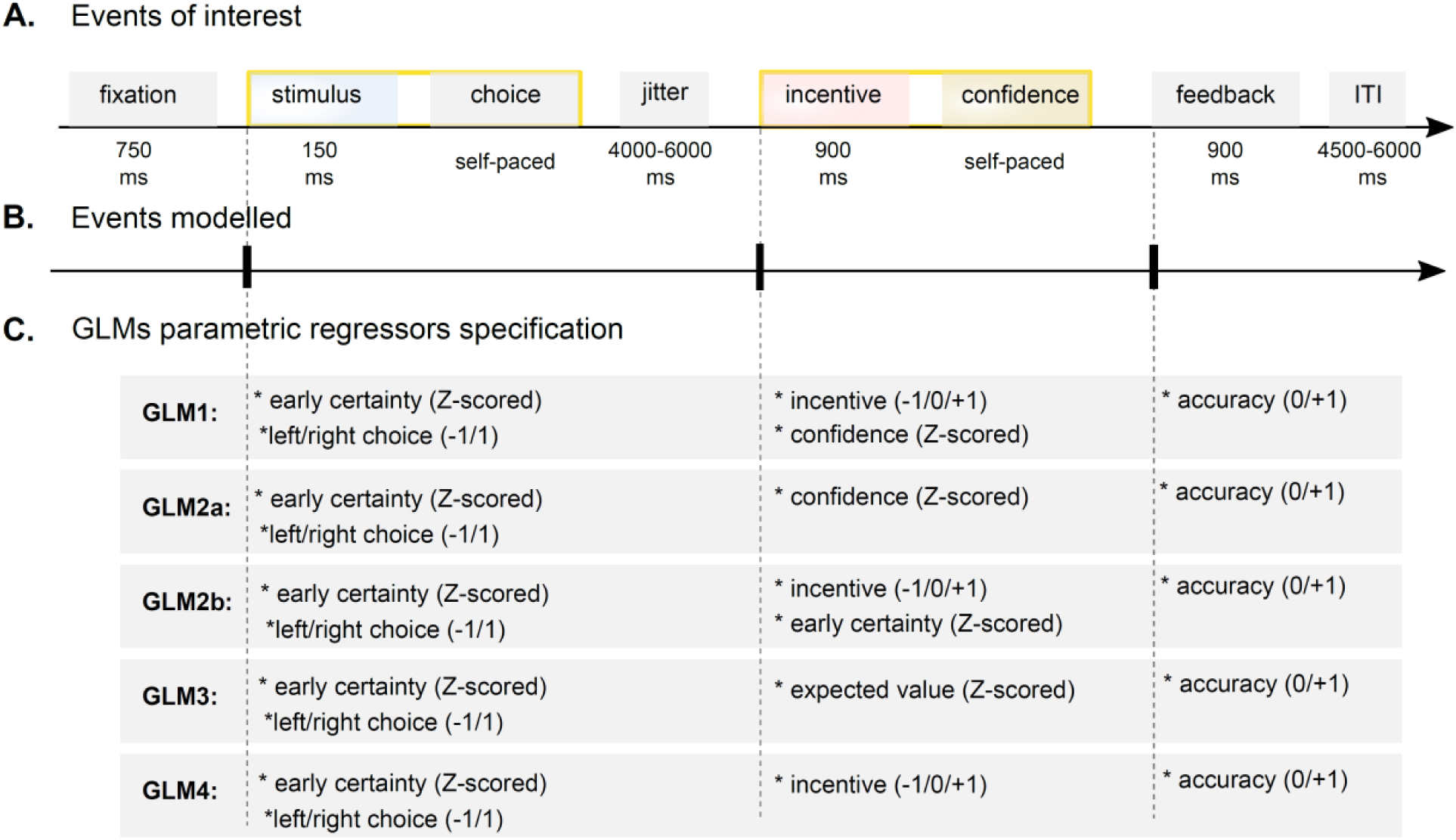
**A-B**. Events of interest. The timeline depicts the succession of events within a trial **A**. Yellow boxes highlight the two events/timing of interest (stimulus/choice and incentive/confidence), that are modelled as stick function for the fMRI analysis. We also modelled the feedback event as a stick function. **C**. GLMs parametric regressors specification. The graph displays the different combination of parametric modulators of each event of interest for all GLMs used to analyze the fMRI data.

#### BOLD signal in the VMPFC correlates significantly with early certainty and incentives but weakly with confidence

Our original hypothesis proposes that incentives bias confidence because those two variables are represented in the same brain area – presumably the VMPFC (De Martino et al., 2013; Lebreton et al., 2015). To test this hypothesis, we built a first fMRI GLM (GLM1) which models 1) early certainty during stimulus and choice (see **Methods** for details), and 2) both incentives and confidence ratings during incentive/rating (**Figure 2C**).

During choice, early certainty positively correlated with activation in the VMPFC, as well as posterior cingulate cortex (PCC) (**Figure 3A**). This replicates several studies that have reported an early and automatic encoding of confidence in the VMPFC (De Martino et al., 2017; Lebreton et al., 2015; Shapiro and Grafton, 2020). Negative correlations were observed in a widespread network of bilateral dorsolateral prefrontal cortex (DLPFC) and rostro-lateral prefrontal cortex (RLPFC), bilateral anterior insula, right putamen, right inferior frontal gyrus, supplementary motor area (SMA), mid- and anterior cingulate cortex and bilateral inferior parietal lobe. This large network has already been implicated in uncertainty and metacognition (Vaccaro and Fleming, 2018).

**Figure 3.**
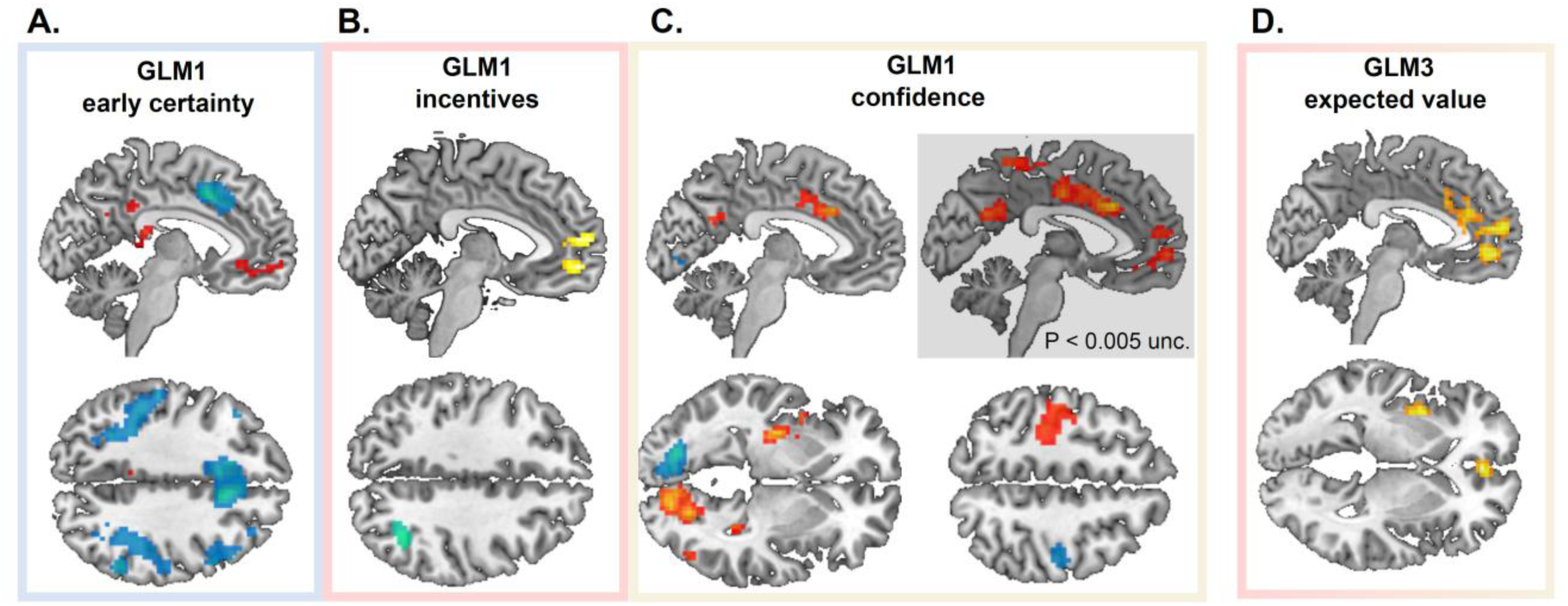
**A-C:** Whole brain statistical BOLD activity correlating with GLM1 early certainty (**A**), incentives (**B**) and confidence (**C**). **D**. Whole brain statistical maps of BOLD activity correlating with GLM3 expected value. Unless otherwise specified, all displayed cluster survived P<0.05 FWE cluster correction. Voxel-wise cluster-defining threshold was set at P < 0.001, uncorrected. Red/yellow clusters: positive activations. Blue clusters: negative activations. For whole-brain activation tables see Supplementary Materials.

During the incentive/rating moment, we found positive correlations between incentive value and activity in the VMPFC - extending to clusters in the dorsomedial prefrontal cortex (**Figure 3B**). This is in line with our hypothesis and with a large neuro-economics literature (Bartra et al., 2013). A small cluster was detected in the occipital lobe, which negatively correlated with incentives.

Finally, regarding subjective confidence, we found significant positive effects in a large, lateralized visuo-motor network including the left primary motor cortex, left putamen and left para-hippocampal gyrus, as well as right cerebellum and right visual cortex (**Figure 3C**). All those activations were mirrored in the negative correlation with confidence (although with lower – and sometimes subthreshold – significance), suggesting these are simply part of the visuo-motor network that process the movement of the cursor on the rating scale - remember that movements of the cursor are operationalized with the left (resp. right) index finger to move the cursor toward the left (resp. right).

Outside those visuo-motor areas, activity in a large cluster in the dorsal anterior cingulate cortex (dACC) and the mid cingulate cortex (MCC) was found to *positively* correlate with confidence. Interestingly, an adjacent region of the dACC *negatively* correlated with early certainty in the choice period (**Figure 3A**). To our surprise, and in contradiction with our hypothesis, no whole-brain significant cluster was found in the VMPFC at our a priori defined statistical threshold. There were, however signs of sub-threshold activations (**Figure 3C**).

#### Accounting for incentive bias in confidence does not restore VMPFC confidence activations

Next, we attempted to understand why no strong correlations with confidence were found in the VMPFC, while the same region robustly encoded early certainty and incentives - two precursors of confidence. We reasoned that because confidence is biased by incentive, shared variance between those two variables could decrease our chances to reveal clear confidence signals during confidence ratings. We therefore built two control GLMs, which differed in how we modelled the incentive/rating period (**Figure 2C**): GLM2a only included confidence as a parametric modulator, while GLM2b included incentive and early certainty (i.e. the precursor of confidence, devoid of incentive shared variance). We defined an anatomical VMPFC ROI (see **Methods** and **Figure 4A**), and extracted individual standardized regression coefficients (t-values) corresponding to the confidence variable in those 3 GLMs (GLM1, GLM2a, GLM2b) (see **Methods**). We then tested whether the difference in the GLM specifications had an impact on these activations at the rating period (GLM1 and 2a: confidence; GLM2b: certainty), using repeated measure ANOVAs. Results showed that activations for GLM2a-confidence and GLM2b-early certainty during incentive/rating period are indistinguishable from GLM1-confidence (ANOVA, main effect of GLMs: F(2,29) = 0.68; P = 0.509), falsifying the hypothesis that the weak confidence activations in VMPFC observed with GLM1 were due to an ill-specified GLM.

**Figure 4.**
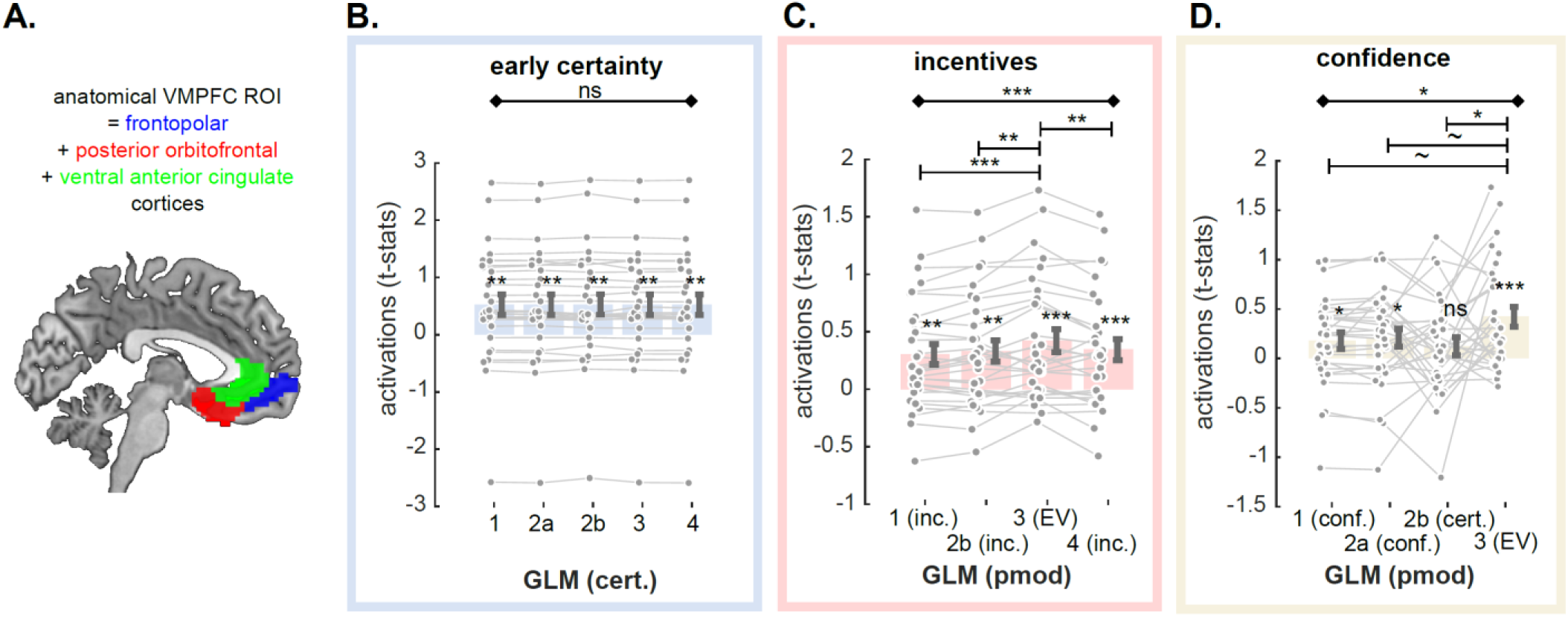
**A**. Anatomical vmPFC region of interest (ROI). **B-D** Comparison of vmPFC activations to different specifications of early certainty during choice moment (B), incentives during incentive/rating moment (C) and confidence during incentive/rating moment (D), as implemented in the different GLMs. Dots represent individual activations; bar and error bars indicate sample mean ± sem. Grey lines highlight within subject variation across the different specifications. Cert: early certainty; Inc.: incentives; conf.: confidence; EV: expected value; Diamond-ended horizontal bars indicate the results of repeated-measure ANOVAs. Dash-ended horizontal bars indicate the result of post-hoc paired t-tests. ∼ P < 0.10; * P<0.05; ** P<0.01; *** P < 0.001

#### BOLD signal in the VMPFC strongly correlates with expected value

Having established that BOLD activity in the VMPFC only weakly correlates with confidence after the incentive display, we proposed a new hypothesis – namely that the VMPFC encodes a signal commensurate to an expected value. The rationale is twofold: first, because confidence represents a subjective probability of being correct, this can be combined with the performance bonus once this is revealed, thereby mechanically generating the expected value (EV) variable. Second, activity in the VMPFC has been repeatedly shown to correlate with EV in different contexts (lotteries, etc.) (Gläscher et al., 2009; Hare et al., 2008; Knutson et al., 2005; McNamee et al., 2013). To test this hypothesis, we built another fMRI GLM similar to the previous ones, but that modeled EV at the time of incentive/rating (GLM 3; see **Figure 2C**).

Whole-brain results showed massive positive correlations between EV and signal in the VMPFC - stretching into anterior medial prefrontal cortex, as well as the ventral and dorsal part of the anterior cingulate cortex and the mid cingulate cortex (**Figure 3D**). There were no clusters with activity negatively related to EV.

#### BOLD signal in the VMPFC correlates better with expected value than with other variables

Although these results seem to validate our second hypothesis, the fact that we observed more activations (wider cluster, lower P-values) at the whole-brain level for EV than for confidence does not constitute a formal statistical test that VMPFC signals might rather correlate with EV than with confidence. Also, they may be caused by the fact that incentives and EV are highly correlated – in other words, VMPFC activations to EV could simply be a result of VMPFC activations to incentives. In order to alleviate those concerns, we built an additional GLM (GLM4), which only included incentive at the incentive/rating period (**Figure 2C**). Again, we extracted VMPFC individual standardized regression coefficients (t-values) corresponding to the early certainty, the incentive and the confidence-related activations in all available GLMs. We tested whether the different specifications had an impact on those activations using repeated measure ANOVAs, and post-hoc t-tests (**Figure 4, Table 1**). Although activations for early certainty during choice moment were similar for all GLMs (ANOVA, main effect of GLM; F(4,29) = 0.24, P = 0.916; **Figure 4B**), GLM specification had an impact on both the incentive activations (ANOVA, main effect of GLM; F(3,29) = 10.67, P = 4.837×10^−6^; **Figure 4C**) and the confidence activations (ANOVA, main effect of GLM; F(3,29) = 3.22, P = 0.027; **Figure 4C**) during incentive/rating moment. In both cases, post-hoc t-tests showed that T-values extracted from the GLM3 that related to the expected value regressor were significantly higher than from other GLMs with a different coding of incentives (GLM1 vs GLM3: t_29_ = 3.90, P = 5.306×10^−4^; GLM2b vs GLM3: t_29_ = 3.38, P = 0.002, GLM4 vs GLM3: t_29_ = 2.97, P = 0.006), and marginally higher from other GLMs with a different coding of confidence (GLM1 vs. GLM3: t_29_ = 1.92, P = 0.064; GLM2a vs. GLM3: t_29_ = 1.72, P = 0.096; GLM2b vs. GLM3: t_29_ = 2.36, P = 0.025). Overall these analyses suggest that the VMPFC combines incentive and confidence signals in the form of an expected value signal.

**Table 1.** Comparison of VMPFC parametric activity (t-values) as a function of model specification (GLMs) The table reports descriptive and inferential statistics on VMPFC ROI parametric activations with three different variables of interest: early certainty effects at choice moment, incentive effects at rating moment and confidence effects at rating moment (see **Fig. 4**). Per effect of interest, results of one-sample t-tests against zero, repeated-measure (RM) ANOVAs on the main effect of GLMS, and post-hoc t-test results are shown.

|  |  |  |  |  |  |
| --- | --- | --- | --- | --- | --- |
| Early certainty | **GLM1** | **GLM2a** | **GLM2b** | **GLM3** | **GLM4** |
|  | 0.52 ± 0.18<br>t<sub>29</sub> = 2.92<br>P = 0.007 | 0.52 ± 0.18<br>t<sub>29</sub> = 2.91<br>P = 0.007 | 0.53 ± 0.18<br>t<sub>29</sub> = 2.93<br>P = 0.007 | 0.53 ± 0.18<br>t<sub>29</sub> = 2.93<br>P = 0.007 | 0.52 ± 0.18<br>t<sub>29</sub> = 2.90<br>P = 0.007 |
|  | **RM ANOVA** | - | - | - | - |
|  | F(4,29) = 0.24<br>P = 0.916 | - | - | -- |  |
| Incentive |  | **GLM1** | **GLM2b** | **GLM3** | **GLM4** |
|  |  | 0.30 ± 0.09<br>t<sub>29</sub> = 3.45<br>P = 0.002 | 0.33 ± 0.09<br>t<sub>29</sub> = 3.60<br>P = 0.001 | 0.42 ± 0.10<br>t<sub>29</sub> = 4.26<br>P = 1.981×10<sup>-4</sup> | 0.34 ± 0.09<br>t<sub>29</sub> = 3.68<br>P = 9.433×10<sup>-4</sup> |
|  | **RM ANOVA** | **T-Test**<br>**[3 vs 1]** | **T-Test**<br>**[3 vs 2b]** | - | **T-Test**<br>**[3 vs 4]** |
|  | F(3,29) = 10.67<br>P = 4.837×10<sup>-6</sup> | 0.12± 0.03<br>t<sub>29</sub> = 3.90<br>P = 5.306×10<sup>-4</sup> | 0.09 ± 0.03<br>t<sub>29</sub> = 3.38<br>P = 0.002 | -- | 0.08 ± 0.03<br>t<sub>29</sub> = 2.97<br>P = 0.006 |
| Confidence |  | **GLM1** | **GLM2a** | **GLM2b** | **GLM3** |
|  |  | 0.18± 0.08<br>t<sub>29</sub> = 2.14<br>P = 0.041 | 0.21 ± 0.09<br>t<sub>29</sub> = 2.30<br>P = 0.028 | 0.12 ± 0.09<br>t<sub>29</sub> = 1.35<br>P = 0.187 | 0.42 ± 0.10<br>t<sub>29</sub> = 4.26<br>P = 1.981×10<sup>-4</sup> |
|  | **RM ANOVA** | **T-Test**<br>**[3 vs 1]** | **T-Test**<br>**[3 vs 2a]** | **T-Test**<br>**[3 vs 2b]** | - |
|  | F(3,29) = 3.22<br>P = 0.027 | 0.24 ± 0.13<br>t<sub>29</sub> = 1.92<br>P = 0.064 | 0.21 ± 0.12<br>t<sub>29</sub> = -1.72<br>P = 0.096 | 0.30 ± 0.13<br>t<sub>29</sub> = 2.36<br>P = 0.025 |  |

#### Qualitative falsification of the EV model of VMPFC activity

Lastly, in order to confirm the conclusions drawn from our quantitative comparison of VMPFC activations, we ran a qualitative falsification exercise (Palminteri et al., 2017). Basically, leveraging the factorial design of our experiment, we can draw qualitative patterns of activations that would be expected under different hypotheses underlying VMPFC activation (**Figure 5A**).

**Figure 5.**
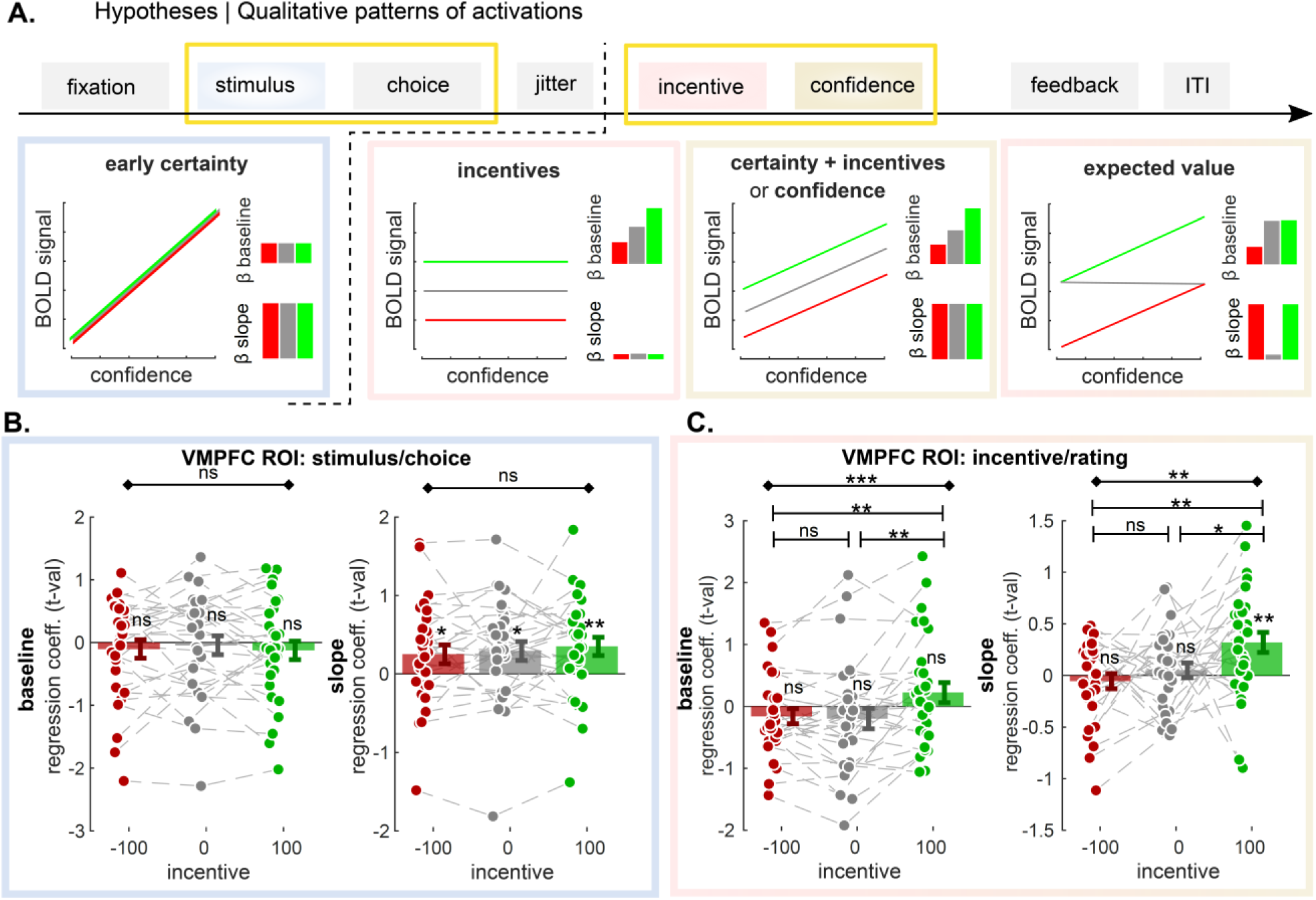
**A**. Qualitative vmPFC activation patterns predicted under different models. The different boxes present how BOLD signal should vary with increasing confidence in the three incentive conditions (green: +100; grey: 0; red: −100), under different hypotheses (i.e. encoding different variables), at different time points. Bar graphs in insets summarize these relationships as expected intercepts (or baseline – top) and slope (bottom). **B-C**. vmPFC ROI analysis. T-values corresponding to baseline and regression slope were extracted in the three incentive conditions, and at the two time-points of interest (B: stimulus/choice; C: incentive/rating). Dots represent individual activations; bar and error bars indicate sample mean ± sem. Grey lines highlight within subject variation across the different incentive conditions. Diamond-ended horizontal bars indicate the results of repeated-measure ANOVAs. Dash-ended horizontal bars indicate the result of post-hoc paired t-tests. ∼ P < 0.10; * P<0.05; ** P<0.01; *** P < 0.001

We therefore designed a last GLM (GLM5) that splits up the task in two time points (stimulus/choice and incentive/rating), and three incentive conditions, and that incorporates a baseline and a regression slope with confidence judgment for all these events. We then extracted the VMPFC activations for all these regressors using our ROI, and compared them with the theorized qualitative patterns we would expect if VMPFC encodes one of these variables (**Figure 5B-C** and **Table 3**). As expected, at the moment of the stimulus/choice, there was no effect of incentive conditions on VMPFC baseline activity, nor on its correlation with confidence – “slope” (ANOVA baseline: F(2,29) = 0.36, P = 0.701; ANOVA correlation with confidence: F(2,29) = 0.56, P = 0.576). Basically, the slopes were significantly positive in all three incentive conditions (Loss: t_29_ = 2.10, P = 0.045; Neutral: t_29_ = 2.43, P = 0.021; Gain: t_29_ = 3.04, P = 0.005), confirming that the VMPFC encodes an early certainty signal.

**Table 2.** Comparison of VMPFC activity at choice moment (t-values), as a function of incentive condition. The table reports descriptive and inferential statistics on VMPFC ROI parametric activations in our three incentive conditions during choice moment, for both baseline activity as well as the correlation with early certainty (i.e. slope) (see **Figure 5B**). Results of RM ANOVAs and one-sample t-tests against 0 are shown.

| Choice/Stim | baseline | Inc. -100 | Inc. 0 | Inc. +100 | RM ANOVA |
| --- | --- | --- | --- | --- | --- |
|  |  | -0.10 ± 0.15<br>t<sub>29</sub> = -0.70<br>P = 0.490 | -0.04 ± 0.15<br>t<sub>29</sub> = -0.30<br>P = 0.770 | -0.13 ± 0.15<br>t<sub>29</sub> = -0.85<br>P = 0.400 | F(2,29) = 0.36<br>P = 0.701 |
|  | slope | Inc. -100 | Inc 0 | Inc. +100 | RM ANOVA |
|  |  | 0.250 ± 0.12<br>t<sub>29</sub> = 2.10<br>P = 0.045 | 0.29 ± 0.12<br>t<sub>29</sub> = 2.43<br>P = 0.021 | 0.35 ± 0.12<br>t<sub>29</sub> = 3.04<br>P = 0.005 | F(2,29) = 0.56<br>P = 0.576 |

**Table 3.**
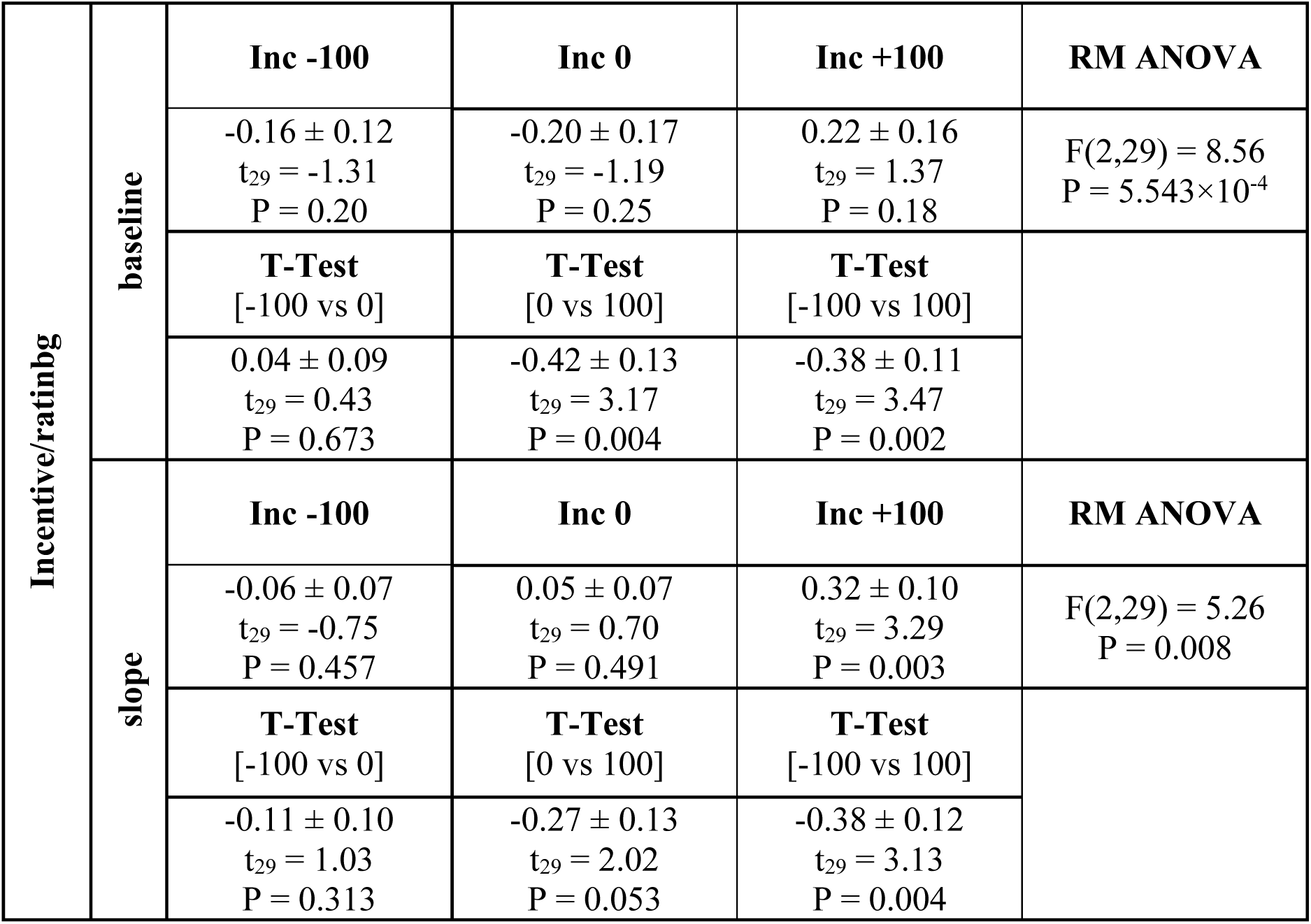
Comparison of VMPFC activity at rating moment (t-values), as a function of incentive condition. The table reports descriptive and inferential statistics on VMPFC ROI parametric activations in our three incentive conditions during rating moment, for both baseline activity as well as the correlation with confidence (i.e. slope) (see **Figure 5C**). Results of one-sample t-tests against 0, RM ANOVAs and post-hoc t-tests are shown.

At rating moment, incentive conditions had an effect on both VMPFC baseline activity, and on its correlation with confidence (ANOVA baseline: F(2,29)= 8.56, P = 5.543×10^−4^; ANOVA correlation with confidence: F(2,29)= 5.26, P = 0.008). Post-hoc testing revealed that VMPFC baseline activity was significantly larger in gain versus loss (t_29_ = 3.47, P = 0.002) and in gain vs neutral conditions (t_29_ = 3.17, P = 0.004), but not in neutral vs loss condition (t_29_ = 0.43, P = 0.673) (see **Table 3**). This constitutes a deviation from a standard linear model of incentives, and suggest that different regions might process incentives in gains and loss contexts (Palminteri and Pessiglione, 2017).

Moreover, we found that the correlation of VMPFC activity with confidence is mostly positively significant in the gain condition (t_29_ = 3.29, P = 0.003), and not in the loss (t_29_ = −0.75, P = 0.457) nor neutral (t_29_ = 0.70, P = 0.491) conditions. The correlation with confidence was therefore significantly higher in gain versus loss (t_29_ = 3.13, P = 0.004) and in gain versus neutral conditions (t_29_ = 2.02, P = 0.053), but not in neutral versus loss condition (t_29_ = 1.03, P = 0.313). While the absence of correlation in the neutral condition would be expected, should the VMPFC encode EV, the lack of correlation in the loss condition was not featured in any model prediction (**Figure 5A**).

Overall, these results explain why EV appears, at first sight, as a better model of VMPFC activation than confidence and/or incentive (correct pattern in gains and neutral conditions), but ultimately falsify this account (absence of positive correlation between VMPFC activation and confidence in the loss condition).

## Discussion

In this study, we set out to investigate the neural signature of incentive bias on confidence estimations, using an fMRI-optimized version of an incentivized perceptual decision-making task (Lebreton et al., 2018). First, at the behavioral level, we replicated the biasing effect of incentives on confidence estimation, in the form of higher confidence in gain contexts and lower confidence in loss context – despite equal difficulty and performance. This result constitutes the fourth independent replication of this bias, initially revealed in perceptual decision making and later generalized in a reinforcement-learning task (Lebreton et al., 2018, 2019a; Ting et al., 2020). Note however, that the bias effect-size remain small – a few average confidence percentage points at the population level –, what sets clear bounds *a priori* on our ability to dissect its precise neurophysiological bases with current (correlational) functional neuroimaging techniques.

Our initial goal and hypothesis was therefore quite simple and modest. In the literature, it is now well established that BOLD signal in the VMPFC correlates with confidence and/or values in a variety of tasks (De Martino et al., 2013, 2017; Lebreton et al., 2015; Lopez-Persem et al., 2020; Morales et al., 2018; Shapiro and Grafton, 2020). We reasoned that if we could unequivocally evidence the co-existence of incentive and confidence signals in VMPFC activity in our task, this would reinforce the intuition that this region plays a role in the observed behavioral phenomenon - the incentive bias on confidence. Our neuroimaging predictions were that 1) VMPFC should automatically correlate with early certainty before and during choice, regardless of the context, and 2) VMPFC should integrate confidence and incentive after the choice and the revealing of the incentive condition. Our broader, speculative neural hypothesis was that during this last confidence judgment step, a third-party metacognitive region or network would sample signal in the VMPFC (Meyniel et al., 2015; Shekhar and Rahnev, 2018), and incidentally end up with a biased confidence estimate incorporating incentive signal.

Our functional MRI investigation of the neural correlates of early certainty confirms our first prediction: BOLD activity in the VMPFC positively correlates with this quantity in all conditions. This result replicates and extends previous studies demonstrating this area to be involved in the initial and automatic processing of confidence during choice (De Martino et al., 2013; Lebreton et al., 2015; Shapiro and Grafton, 2020). In parallel with this positive correlation in the VMPFC, we also observed wide-spread negative correlations in the DLPFC, DMPFC and insula, a network robustly associated with both metacognition and uncertainty (Molenberghs et al., 2016; Morales et al., 2018; Vaccaro and Fleming, 2018). Contrary to our second prediction, we only found weak evidence (i.e. at a lower statistical threshold than the one we defined *a priori*) for confidence encoding in the VMPFC. Robust activations were nonetheless observed in the dACC, a region known to be recruited in metacognitive judgments (Bang and Fleming, 2018; Fleming et al., 2012).

Given that the lack of robust confidence signal in the VMPFC is somewhat in contradiction with what we expected from our previous work, as well as numerous other reports in the literature (De Martino et al., 2013, 2017; Lebreton et al., 2015; Lopez-Persem et al., 2020; Morales et al., 2018; Shapiro and Grafton, 2020), we formulated an alternative hypothesis: we proposed that VMPFC could encode a signal commensurate to an expected reward (or expected value EV), i.e. incorporating the subjective probability of being correct with the potential incentive bonus when revealed. Whole-brain activations and ROI quantitative analyses clearly showed that this second hypothesis seems to give a better account of VMPFC BOLD activations. Expected value signals are frequently reported in the VMPFC, but mostly in reinforcement-learning contexts, where they are critical to both choices between available options and learning – i.e. value updating, through the computation of prediction errors (Chase et al., 2015). In the present perceptual task, there is no learning, therefore no explicit need to encode EV.

Because quantitative comparisons of hypotheses are notoriously hard to interpret, we decided to leverage the factorial aspect of our design to proceed to a qualitative hypothesis falsification, to validate – or falsify – the EV account of VMPFC activity (Palminteri et al., 2017). In short, different hypotheses about what should be contained in VMPFC signal (EV, confidence and/or incentives) predict different patterns of activations (baseline and correlation with confidence) in our different incentive conditions. From activity extracted in an anatomical VMPFC ROI, it is clear that activity in the VMPFC correlates with confidence only in the gain context, once the incentive has been revealed. This finding explains why the EV hypothesis gets more quantitative support than the confidence and/or incentives hypotheses (as the VMPFC activity pattern is similar to the EV predictions in the gain and neutral context). However, it also ultimately falsifies this EV hypothesis as well (as VMPFC activity does not seem to correlate with confidence in the loss context). Interestingly VMPFC did correlate with early certainty – a precursor of confidence - in all conditions before the incentives are revealed. Therefore, it does not seem that VMPC fails to activate in the neutral and loss conditions, but rather that the signal is actively suppressed once those contexts are explicit. In summary, we believe that our results show a complex picture of disruptions of confidence signals within the VMPFC in response to motivational signals.

The notion that there are different brain networks which execute symmetric computations in gains vs. loss contexts is increasingly popular (Palminteri and Pessiglione, 2017; Seymour et al., 2015). In the present dataset we did not find any region correlating either positively or negatively with confidence in the loss context, even when exploring at the whole brain level with very lenient statistical thresholds. One promising area, that has repeatedly been found to be responsive to loss anticipation and that correlated positively with subjective confidence in our data, is the dACC. However, when we performed a similar falsification exercise within the dACC as we used within the VMPFC (see **Suppl. Mat**.), the results were similar to the VMPFC activation patterns: dACC activity only correlated with confidence within the gain contexts. In summary, the neurobiological bases of confidence judgments in loss contexts remain an open question.

Our results have important implications, as the influences of motivational processes on confidence could lead to dysfunctional behaviors. Indeed, several psychiatric disorders such as addiction, obsessive-compulsive disorder and schizophrenia have been associated with disrupted incentive processing, and studies have additionally demonstrated distorted confidence estimations in these groups (Hoven et al., 2019). Moreover, the regions involved in incentive processing and confidence estimation such as the VMPFC, VS, insula and ACC are all consistently implicated in these disorders (Chai et al., 2011; Goldstein and Volkow, 2011; Namkung et al., 2017; Thorsen et al., 2018). Therefore, the relation between motivational processes and confidence estimation and their role in psychopathology warrants future investigation.

In conclusion, we show that although the VMPFC superficially seems to encode both value and metacognitive signals, these metacognitive signals are only present during the prospect of gain, but are disrupted in a context with loss or no monetary prospects. Studies targeting this problem within a finer spatial (Kepecs and Mainen, 2012; Lopez-Persem et al., 2020; Middlebrooks et al., 2013), as well as temporal scale (Desender et al., 2016), could help with resolving and better comprehending biased confidence judgments and metacognition overall.

## Methods

### Participants

We included thirty-three right-handed healthy participants with normal or corrected to normal vision. Exclusion criteria were an IQ below 80, insufficient command of the Dutch language or MRI contraindications. All experimental procedures were approved by the Medical Ethics Committee of the Academic Medical Center, University of Amsterdam (METC 2015_319) and participants gave written informed consent. Participants were compensated with a base amount of €40 and additional gains based on task performance. Session-level behavioral and fMRI data was excluded when task accuracy was below 60% or when subjects did not show sufficient variation in their confidence reports (standard deviation of confidence judgments <5 confidence points), and session-level fMRI data when participants displayed more than 3.5 mm head movement. This led to the inclusion of 32 participants for the behavioral analyses and 30 for the fMRI analyses, of which four participants contributed only one of two task sessions.

### Decision-making and confidence judgment task

We adapted the task from Lebreton et al. (2018) for use in an fMRI environment with fMRI suitable timing intervals. For an overview and details, see **Figure 1A**. All tasks used in this study were implemented using MATLAB® (MathWorks Inc., Sherborn, MA, USA) and the COGENT toolbox (www.vislab.ucl.ac.uk/cogent.php).

### Study procedure

On the day of testing, subjects were first assessed for clinical and demographic data, then performed a ten-trial practice session, once outside and once inside the scanner, to get acquainted with the task. Before performing the main task, subjects were instructed that it was important to be as accurate as possible as this would maximize their earnings, but to also give accurate confidence judgments. All subjects initially performed a 144-trial calibration session inside the scanner to tailor the difficulty levels of the task to each individual and to keep performance constant across subjects - see Lebreton et al. (2018) for details.

Two sessions of the main task were performed in the fMRI scanner, each consisting of 72 trials with 24 trials per incentive condition, presented in a random order. The practice task, calibration and main sessions were projected onto an Iiyama monitor in the fMRI environment, which subjects could see through a 45-degree angle mirror fixed to the head coil. After completing the fMRI task, six random trials were drawn (i.e. two of each incentive condition) on which the payment was based. If subjects made an accurate choice, they would either gain or avoid losing points, whereas they would miss out on gaining or lose points when making an error. In the neutral trials, nothing was at stake. Finally, the total amount of points was converted to money.

### Behavioral Measures

We extracted various trial-by-trial experimental factors (evidence, incentive and difficulty level) and behavioral measures (accuracy, subjective confidence ratings, reaction times). Two additional variables were computed as combinations of those experimental factors and behavioral measures: early certainty and expected value.

#### Early certainty

We built an ‘early certainty’ variable to analyze brain activity at choice moment, when the brain possibly automatically encodes a confidence signal that is not yet biased by incentives (Lebreton et al., 2015). We reasoned that such a signal should be highly correlated with confidence, while exhibiting no statistical relationships with incentives. Therefore, we used a leave-one-trial-out approach to obtain trial-by-trial estimations of early certainty (Bang and Fleming, 2018). We fitted a generalized linear regression model to each subject’s subjective confidence ratings using choice and stimulus features as predictors (i.e. log-transformed reaction times, evidence, accuracy and the interaction between accuracy and evidence), using the whole individual dataset but trial *X*. We then applied this model’s beta estimates to generate predictions about the early certainty in trial *X*, using the choice and stimulus features of trial *X*. This process was repeated for every trial, resulting in a trial-by-trial prediction of early certainty based on stimulus features at choice moment. The resulting early certainty signal featured high correlation with confidence, and no statistical relationship with incentives (see **Suppl. Mat**. for more details).

#### Expected value

we computed a value-based measure of expected value (EV). In our task paradigm, EV was computed as an integrative signal of early certainty (i.e. the non-biased probability of being correct) and the incentive value (i.e. value-context of the current trial). Early certainty ratings represent the subjects’ probability of being correct, and thus the probability of gaining (or avoid losing) the incentive at stake. Thus, EV corresponds to 0 in the neutral condition (no value is expected to be gained or lost), is equal to early certainty in the gain condition (e.g. being 100% certain results in a maximal EV in a positive incentive environment), and is equal to early certainty – 100 (e.g. being 100% certain in a loss trial results in an EV of 0, as you avoid losing).

### Behavioral Analyses

All behavioral analyses were performed using MATLAB and the R environment (RStudio Team (2015). RStudio: Integrated Development for R. RStudio, Inc., Boston, MA). For the statistical analyses reported in the main text, we used linear mixed-effects models (estimated with the fitglme function in MATLAB) to model accuracy, reaction times and confidence. For all three trial-by-trial dependent variables we used the absolute incentive value (|V|) and the net incentive value (V) as predictor variables. All mixed models included random intercepts and random slopes. Additional control analyses are reported in the **Suppl. Mat**.

### fMRI acquisition & preprocessing

fMRI data was acquired by using a 3.0 Tesla Intera MRI scanner (Philips Medical Systems, Best, The Netherlands). Following the acquisition of a T1-weighted structural anatomical image, 37 axial T2*-weighted EPI functional slices sensitive to blood oxygenation level-dependent (BOLD) contrast were acquired. A multi echo (3 echoes) combine interleaved scan sequence was applied, designed to optimize functional sensitivity in all parts of the brain (Poser et al., 2006). The following imaging parameters were used: TR, 2.375 seconds; TEs, 9.0ms, 24.0ms, and 43.8ms, (total echo train length: 75ms); 3 mm (isometric) voxel size; 37 transverse slices; 3 mm slice thickness; 0.3 mm slice-gap. Two experimental sessions were carried out, each consisting of 570 volumes. All further analyses were performed using MATLAB® (MathWorks Inc., Sherborn, MA, USA) with SPM12 software (Wellcome Department of Cognitive Neurology, London, UK).

Raw multi-echo functional scans were weighed and combined into 570 volumes per scan session. During the combining process, realignment was performed on the functional data by using linear interpolation to the first volume. The first 30 dummy scans were discarded. The remaining functional images were co-registered with the T1-weighted structural image, segmented for normalization to MNI space and smoothed by using a Gaussian kernel of 6 mm at full-width at half-maximum.

Due to sudden motion, in combination with the interleaved scanning method, a number of subjects showed artifacts in some functional volumes. In order to reduce those artifacts, the Art-Repair toolbox (Mazaika et al., 2007) was used to detect large volume-to-volume movement and repair outlier volumes. The toolbox identifies outliers by using a threshold for the variation of the mean intensity of the BOLD signal and a volume-to-volume motion threshold. A threshold of 1.5% variation from the mean intensity was used to detect and repair volume outliers by interpolating from the adjacent volumes (n=12).

### fMRI Analyses

All fMRI analyses were conducted using SPM12. All general linear models (GLMs) were estimated on subject-level with two moments of interest: the moment of choice (i.e. presentation of the Gabor patches) and the moment of incentive presentation/confidence rating (**Fig 2**). The rating moment follows the presentation of the incentive after 900 ms, hence the decision to analyze them as a single moment of interest. Moreover, the GLMs also included a regressor for the feedback moment, which was not of interest for analysis, but was intended to explain variance in neural responses related to value and accuracy feedback, but unrelated to the decision-making process.

When using parametric modulators in our GLMs, those were not orthogonalized and competed to explain variance. Nuisance regressors consisting of 6 motion parameters were included in all GLMs. Regressors were modeled separately for each scan session and constants were included to account for between-session differences in mean activation. All events were modeled by convolving a series of delta functions with the canonical HRF at the onset of each event and were linearly regressed onto the functional BOLD-response signal. Low frequency noise was filtered with a high pass filter with a cut off of 128 seconds. All contrasts were computed at subject level and taken to a group level mixed effect analysis using one-sample t-tests.

We controlled for the number of sessions while making the first-level contrasts. We assessed group-level main effects by applying one-sample t-tests against 0 to these contrast images. All whole-brain activation maps were thresholded using family-wise error correction for multiple correction (FWE) at cluster level (P_FWE_clu_ < 0.05), with a voxel cluster-defining threshold of P<.001 uncorrected.

#### GLM1: Neural signatures of certainty, incentive and confidence

GLM1 consisted of three regressors for the three moments of interest: ‘choice’, ‘incentive/rating’ and ‘feedback’, to which one or more parametric modulators (pmod) were added (**Fig 2**). The regressors were specified as stick function time-locked to the onset of the events. The choice regressor was modulated by two pmods: early certainty (z-scored before entering the GLM) and button press (left/right choice) in order to control for activity related to motor preparation. The incentive/rating regressor was modulated by two pmods: incentive value and subjective confidence level (z-scored). Lastly, the feedback regressor was modulated by a pmod of accuracy.

#### GLM2a: control for incentive bias 1

GLM2a consists of the same regressors as GLM1, except that rating moment is only modulated by confidence judgments (i.e. we deleted the incentive modulator).

#### GLM2b: control for incentive bias 2

GLM2b consists of the same regressors as GLM1, except that the pmod of confidence judgments at rating moment is replaced by a pmod for early certainty.

#### GLM3: Neural signatures of expected value

GLM3 consists of the same regressors as GLM1, except that rating moment is modulated by a single pmod of EV.

#### GLM4: control for incentive

GLM4 consists of the same regressors as GLM1, except that rating moment is only modulated by incentives (i.e. we deleted the confidence judgment modulator).

#### GLM5: qualitative patterns of activations

GLM5 includes a regressor for all three incentives at two timepoints of interest: choice and rating moment, as well as a regressor at feedback moment. All regressors at choice moment were modulated by a pmod of early certainty and button press (L/R). All regressors at rating moment were modulated by a pmod of confidence judgment. The feedback regressor was modulated by accuracy. This GLM allowed us to investigate activity related to both baseline and the regression slope with early certainty or confidence judgment for these events.

#### Regions of interest

To avoid circular inference, we took an independent anatomical ROI of the VMPFC from the Brainnetome Atlas (Fan et al., 2016). We included three areas along the ventral medial axis for the VMPFC ROI. Using this ROI, we extracted individual t-statistics (i.e. normalized beta estimates – see (Lebreton et al., 2019b) from contrasts of interest, and statistically compared them using paired t-tests or repeated measure ANOVAs.

## References

Allen, M., Frank, D., Schwarzkopf, D.S., Fardo, F., Winston, J.S., Hauser, T.U., and Rees, G. (2016). Unexpected arousal modulates the influence of sensory noise on confidence. ELife 5, e18103.

Bang, D., and Fleming, S.M. (2018). Distinct encoding of decision confidence in human medial prefrontal cortex. Proceedings of the National Academy of Sciences of the United States of America 115, 6082–6087.

Bartra, O., McGuire, J.T., and Kable, J.W. (2013). The valuation system: A coordinate-based metaanalysis of BOLD fMRI experiments examining neural correlates of subjective value. NeuroImage 76, 412–427.

Chai, X.J., Whitfield-Gabrieli, S., Shinn, A.K., Gabrieli, J.D.E., Nieto Castañón, A., McCarthy, J.M., Cohen, B.M., and Öngür, D. (2011). Abnormal Medial Prefrontal Cortex Resting-State Connectivity in Bipolar Disorder and Schizophrenia. Neuropsychopharmacology 36, 2009–2017.

Chase, H.W., Kumar, P., Eickhoff, S.B., and Dombrovski, A.Y. (2015). Reinforcement learning models and their neural correlates: An activation likelihood estimation meta-analysis. Cogn Affect Behav Neurosci 15, 435–459.

Chib, V.S., Rangel, A., Shimojo, S., and O’Doherty, J.P. (2009). Evidence for a Common Representation of Decision Values for Dissimilar Goods in Human Ventromedial Prefrontal Cortex. The Journal of Neuroscience 29, 12315–12320.

Daw, N.D., Niv, Y., and Dayan, P. (2005). Uncertainty-based competition between prefrontal and dorsolateral striatal systems for behavioral control. Nature Neuroscience 8, 1704–1711.

De Martino, B., Fleming, S.M., Garrett, N., and Dolan, R.J. (2013). Confidence in value-based choice. Nat Neurosci 16, 105–110.

De Martino, B., Bobadilla-Suarez, S., Nouguchi, T., Sharot, T., and Love, B.C. (2017). Social Information Is Integrated into Value and Confidence Judgments According to Its Reliability. The Journal of Neuroscience 37, 6066–6074.

Desender, K., Van Opstal, F., Hughes, G., and Van den Bussche, E. (2016). The temporal dynamics of metacognition: Dissociating task-related activity from later metacognitive processes. Neuropsychologia 82, 54–64.

Donoso, M., Collins, A.G.E., and Koechlin, E. (2014). Foundations of human reasoning in the prefrontal cortex. Science 344, 1481–1486.

Fan, L., Li, H., Zhuo, J., Zhang, Y., Wang, J., Chen, L., Yang, Z., Chu, C., Xie, S., Laird, A.R., et al. (2016). The Human Brainnetome Atlas: A New Brain Atlas Based on Connectional Architecture. Cerebral Cortex 26, 3508–3526.

Fleming, S.M., and Daw, N.D. (2017). Self-evaluation of decision-making: A general Bayesian framework for metacognitive computation. Psychological Review 124, 91–114.

Fleming, S.M., and Dolan, R.J. (2012). The neural basis of metacognitive ability. Phil. Trans. R. Soc. B 367, 1338–1349.

Fleming, S.M., Huijgen, J., and Dolan, R.J. (2012). Prefrontal Contributions to Metacognition in Perceptual Decision Making. J Neurosci 32, 6117–6125.

Folke, T., Jacobsen, C., Fleming, S.M., and Martino, B.D. (2016). Explicit representation of confidence informs future value-based decisions. Nature Human Behaviour 1, 0002.

Giardini, F., Coricelli, G., Joffily, M., and Sirigu, A. (2008). Overconfidence in Predictions as an Effect of Desirability Bias. In Advances in Decision Making Under Risk and Uncertainty, P.M. Abdellaoui, and P.D.J.D. Hey, eds. (Springer Berlin Heidelberg), pp. 163–180.

Gläscher, J., Hampton, A.N., and O’Doherty, J.P. (2009). Determining a Role for Ventromedial Prefrontal Cortex in Encoding Action-Based Value Signals During Reward-Related Decision Making. Cereb Cortex 19, 483–495.

Goldstein, R.Z., and Volkow, N.D. (2011). Dysfunction of the prefrontal cortex in addiction: neuroimaging findings and clinical implications. Nature Reviews Neuroscience 12, 652–669.

Haber, S.N., and Behrens, T.E.J. (2014). The Neural Network Underlying Incentive-Based Learning: Implications for Interpreting Circuit Disruptions in Psychiatric Disorders. Neuron 83, 1019–1039.

Haber, S.N., and Knutson, B. (2009). The Reward Circuit: Linking Primate Anatomy and Human Imaging. Neuropsychopharmacology 35, 4–26.

Hare, T.A., O’Doherty, J., Camerer, C.F., Schultz, W., and Rangel, A. (2008). Dissociating the Role of the Orbitofrontal Cortex and the Striatum in the Computation of Goal Values and Prediction Errors. J. Neurosci. 28, 5623–5630.

Heilbron, M., and Meyniel, F. (2019). Confidence resets reveal hierarchical adaptive learning in humans. PLOS Computational Biology 15, e1006972.

Hoven, M., Lebreton, M., Engelmann, J.B., Denys, D., Luigjes, J., and van Holst, R.J. (2019). Abnormalities of confidence in psychiatry: an overview and future perspectives. Translational Psychiatry 9, 1–18.

Jönsson, F.U., Olsson, H., and Olsson, M.J. (2005). Odor emotionality affects the confidence in odor naming. Chemical Senses 30, 29–35.

Kable, J.W., and Glimcher, P.W. (2009). The Neurobiology of Decision: Consensus and Controversy. Neuron 63, 733–745.

Kahnt, T., Heinzle, J., Park, S.Q., and Haynes, J.-D. (2011). Decoding different roles for vmPFC and dlPFC in multi-attribute decision making. NeuroImage 56, 709–715.

Kepecs, A., and Mainen, Z.F. (2012). A computational framework for the study of confidence in humans and animals. Philosophical Transactions of the Royal Society B: Biological Sciences 367, 1322–1337.

Knutson, B., Fong, G.W., Bennett, S.M., Adams, C.M., and Hommer, D. (2003). A region of mesial prefrontal cortex tracks monetarily rewarding outcomes: characterization with rapid event-related fMRI. NeuroImage 18, 263–272.

Knutson, B., Taylor, J., Kaufman, M., Peterson, R., and Glover, G. (2005). Distributed Neural Representation of Expected Value. J. Neurosci. 25, 4806–4812.

Koellinger, P., and Treffers, T. (2015). Joy Leads to Overconfidence, and a Simple Countermeasure. PLOS ONE 10, e0143263.

Lebreton, M., Jorge, S., Michel, V., Thirion, B., and Pessiglione, M. (2009). An Automatic Valuation System in the Human Brain: Evidence from Functional Neuroimaging. Neuron 64, 431–439.

Lebreton, M., Abitbol, R., Daunizeau, J., and Pessiglione, M. (2015). Automatic integration of confidence in the brain valuation signal. Nat Neurosci 18, 1159–1167.

Lebreton, M., Langdon, S., Slieker, M.J., Nooitgedacht, J.S., Goudriaan, A.E., Denys, D., Holst, R.J. van, and Luigjes, J. (2018). Two sides of the same coin: Monetary incentives concurrently improve and bias confidence judgments. Science Advances 4, eaaq0668.

Lebreton, M., Bacily, K., Palminteri, S., and Engelmann, J.B. (2019). tContextual influence on confidence judgments in human reinforcement learning. PLOS Computational Biology 15, e1006973.

Lebreton, M., Bavard, S., Daunizeau, J., and Palminteri, S. (2019). tAssessing inter-individual differences with task-related functional neuroimaging. Nat Hum Behav 3, 897–905.

Levy, D.J., and Glimcher, P.W. (2011). Comparing Apples and Oranges: Using Reward-Specific and Reward-General Subjective Value Representation in the Brain. J. Neurosci. 31, 14693–14707.

Levy, I., Lazzaro, S.C., Rutledge, R.B., and Glimcher, P.W. (2011). Choice from Non-Choice: Predicting Consumer Preferences from Blood Oxygenation Level-Dependent Signals Obtained during Passive Viewing. The Journal of Neuroscience 31, 118–125.

Lopez-Persem, A., Bastin, J., Petton, M., Abitbol, R., Lehongre, K., Adam, C., Navarro, V., Rheims, S., Kahane, P., Domenech, P., et al. (2020). Four core properties of the human brain valuation system demonstrated in intracranial signals. Nature Neuroscience 23, 664–675.

Massoni, S. (2014). Emotion as a boost to metacognition: How worry enhances the quality of confidence. Consciousness and Cognition 29, 189–198.

Mazaika, P., Whitfield-Gabrieli, S., and Reiss, A.L. (2007). Artifact repair of fMRI data from high motion clinical subjects. Neuroimage 36, S142.

McNamee, D., Rangel, A., and O’Doherty, J.P. (2013). Category-dependent and category-independent goal-value codes in human ventromedial prefrontal cortex. Nature Neuroscience 16, 479–485.

Meyniel, F., Sigman, M., and Mainen, Z.F. (2015). Confidence as Bayesian Probability: From Neural Origins to Behavior. Neuron 88, 78–92.

Middlebrooks, P.G., Abzug, Z.M., and Sommer, M.A. (2013). Studying metacognitive processes at the single neuron level. In The Cognitive Neuroscience of Metacognition, (Springer-Verlag Berlin Heidelberg), pp. 225–244.

Molenberghs, P., Trautwein, F.M., Böckler, A., Singer, T., and Kanske, P. (2016). Neural correlates of metacognitive ability and of feeling confident: a large-scale fMRI study. Social Cognitive and Affective Neuroscience 11, 1942–1951.

Morales, J., Lau, H., and Fleming, S.M. (2018). Domain-General and Domain-Specific Patterns of Activity Supporting Metacognition in Human Prefrontal Cortex. J. Neurosci. 38, 3534–3546.

Namkung, H., Kim, S.-H., and Sawa, A. (2017). The Insula: An Underestimated Brain Area in Clinical Neuroscience, Psychiatry, and Neurology. Trends in Neurosciences 40, 200–207.

Padoa-Schioppa, C. (2007). Orbitofrontal Cortex and the Computation of Economic Value. Annals of the New York Academy of Sciences 1121, 232–253.

Padoa-Schioppa, C., and Assad, J.A. (2006). Neurons in the orbitofrontal cortex encode economic value. Nature 441, 223–226.

Palminteri, S., and Pessiglione, M. (2017). Chapter 23 - Opponent Brain Systems for Reward and Punishment Learning: Causal Evidence From Drug and Lesion Studies in Humans. In Decision Neuroscience, J.-C. Dreher, and L. Tremblay, eds. (San Diego: Academic Press), pp. 291–303.

Palminteri, S., Wyart, V., and Koechlin, E. (2017). The Importance of Falsification in Computational Cognitive Modeling. Trends in Cognitive Sciences 21, 425–433.

Pessiglione, M., and Lebreton, M. (2015). From the Reward Circuit to the Valuation System: How the Brain Motivates Behavior. In Handbook of Biobehavioral Approaches to Self-Regulation, G.H.E. Gendolla, M. Tops, and S.L. Koole, eds. (New York, NY: Springer), pp. 157–173.

Plassmann, H., O’Doherty, J., and Rangel, A. (2007). Orbitofrontal Cortex Encodes Willingness to Pay in Everyday Economic Transactions. J. Neurosci. 27, 9984–9988.

Poser, B.A., Versluis, M.J., Hoogduin, J.M., and Norris, D.G. (2006). BOLD contrast sensitivity enhancement and artifact reduction with multiecho EPI: Parallel-acquired inhomogeneity-desensitized fMRI. Magnetic Resonance in Medicine 55, 1227–1235.

Pouget, A., Drugowitsch, J., and Kepecs, A. (2016). Confidence and certainty: distinct probabilistic quantities for different goals. Nat Neurosci 19, 366–374.

Rangel, A., and Hare, T. (2010). Neural computations associated with goal-directed choice. Current Opinion in Neurobiology 20, 262–270.

Seymour, B., Maruyama, M., and De Martino, B. (2015). When is a loss a loss? Excitatory and inhibitory processes in loss-related decision-making. Current Opinion in Behavioral Sciences 5, 122–127.

Shapiro, A.D., and Grafton, S.T. (2020). Subjective value then confidence in human ventromedial prefrontal cortex. PLOS ONE 15, e0225617.

Shekhar, M., and Rahnev, D. (2018). Distinguishing the Roles of Dorsolateral and Anterior PFC in Visual Metacognition. J. Neurosci. 38, 5078–5087.

Thorsen, A.L., Hagland, P., Radua, J., Mataix-Cols, D., Kvale, G., Hansen, B., and van den Heuvel, O.A. (2018). Emotional Processing in Obsessive-Compulsive Disorder: A Systematic Review and Meta-analysis of 25 Functional Neuroimaging Studies. Biological Psychiatry: Cognitive Neuroscience and Neuroimaging 3, 563–571.

Ting, C.-C., Palminteri, S., Engelmann, J.B., and Lebreton, M. (2020). Robust valence-induced biases on motor response and confidence in human reinforcement learning. Cogn Affect Behav Neurosci.

Tremblay, L., and Schultz, W. (1999). Relative reward preference in primate orbitofrontal cortex. Nature 398, 704–708.

Vaccaro, A.G., and Fleming, S.M. (2018). Thinking about thinking: A coordinate-based meta-analysis of neuroimaging studies of metacognitive judgements. Brain and Neuroscience Advances 2, 2398212818810591.

Vinckier, F., Gaillard, R., Palminteri, S., Rigoux, L., Salvador, A., Fornito, A., Adapa, R., Krebs, M.O., Pessiglione, M., and Fletcher, P.C. (2016). Confidence and psychosis: a neuro-computational account of contingency learning disruption by NMDA blockade. Molecular Psychiatry 21, 946–955.

