## Supplementary Materials for "How motivational signals disrupt metacognitive signals in the human VMPFC"

#### Additional Behavioral Analyses: Properties of Confidence Judgments

Similarly to Lebreton et al. (2018), we performed additional behavioral analyses to confirm three main properties of confidence judgements, as theorized in a recent paper by Sanders and colleagues (Sanders et al., 2016). There, the authors outlined three main properties of confidence judgments, which should be observed if participants compute the probability of a choice being correct given some level of noisy evidence: (1) confidence ratings correlate with the probability of being correct; (2) the link between confidence ratings and evidence is positive for correct and negative for incorrect responses; (3) the link between evidence and performance differs between high and low confidence trials.

- To assess the first property, we sorted trials according to the confidence ratings at the individual level. Then, we averaged trials over 8 bins per participant, and computed the frequency of correct choices in each bin. Finally, the correlation between the bins' confidence and performance was computed at the individual level. These measures were positively correlated ($R = 0.59 \pm 0.06$; **Figure S1A**).
- To assess the second property, the following linear regression was estimated at the individual level, using all trials from the confidence elicitation task (Model 1):

  $\text{Conf} = \beta 0 + \beta_1 \times \text{Correct} \times \text{Evidence} + \beta_2 \times \text{Incorrect} \times \text{Evidence}$,

  where **Incorrect** is a dummy variable coding for incorrect answers, and **Correct** is a dummy variable coding for correct answers. Then, we tested the parameters of this model at the population level using one-sample t-tests. The results (**Figure S1B**), summarized in the table below (**Table S1**), demonstrate that confidence judgments are indeed positively associated with evidence for correct trials, and negatively for incorrect trials.
- To assess the third property, we proceeded similarly to the second: the following logistic regression was estimated at the individual level, using all trials (Model 2).

  $\text{Correct} = \beta 0 + \beta_1 \times \text{High} \times \text{Evidence} + \beta_2 \times \text{Low} \times \text{Evidence}$,

  where **High** is a dummy variable coding for high confidence trials (i.e. confidence > median(confidence)), and **Low** is a dummy variable coding for low confidence trials (i.e. confidence ≤ median(confidence)). Then, the parameters of this model were tested at the population level, using one-sample t-tests. The results (**Figure S1C**), summarized in the table below (**Table S1**), indeed demonstrate that the curve has a steeper slope in the high than in the low confidence trials, as was expected.

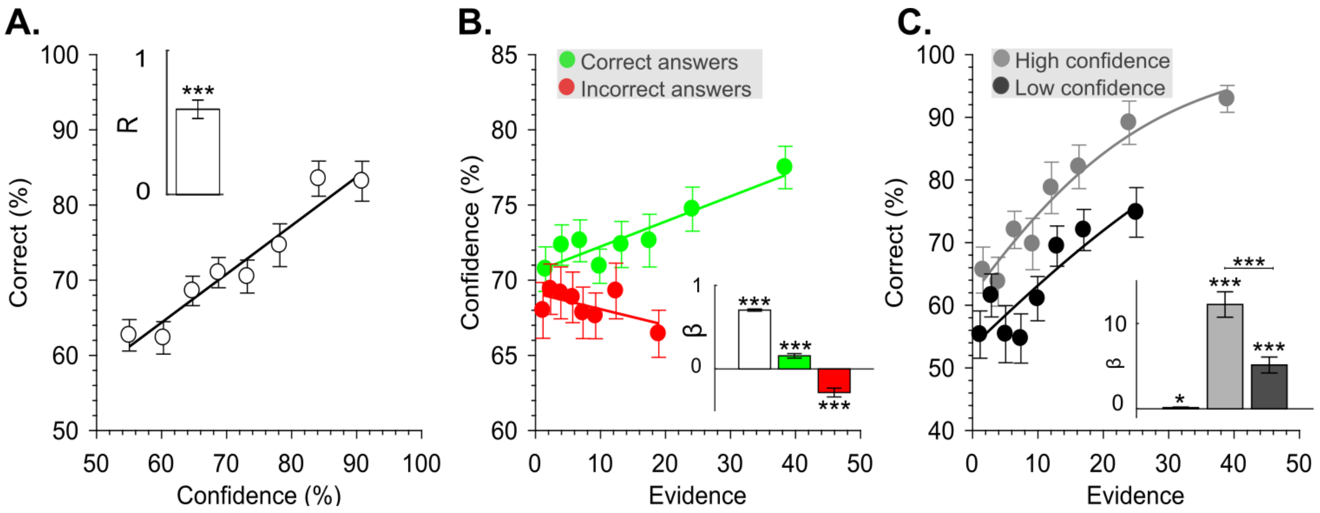

**Figure S1 | Properties of Confidence Judgments**

A: observed performance (% correct choices) as a function of reported confidence. B: reported confidence as function of evidence for correct (green) and incorrect (red) choices. C: observed performance (% correct choices) as a function of evidence, for high (gray) and low (black) confidence trials. The insets presented on the side of each graph depict the results of the population-level analyses on the correlation coefficients (A) or on the regression coefficients (B and C). Error bars indicate inter-subject standard errors of the mean. *: P<0.05; **: P<0.01; ***P<0.001

| Model 1 (Figure S1B) | |
|---|---|
| **Intercept ($\beta_0$)** | $\beta = 0.71 \pm 0.01$<br>$t_{31} = 53.06$<br>$P = 5.353\times10^{-32}$ |
| **Confidence/Evidence Correct Answers ($\beta_1$)** | $\beta = 0.16 \pm 0.03$<br>$t_{31} = 5.86$<br>$P = 1.833\times10^{-6}$ |
| **Confidence/Evidence Incorrect Answers ($\beta_2$)** | $\beta = -0.28 \pm 0.05$<br>$t_{31} = -5.36$<br>$P = 7.703\times10^{-6}$ |
| **Model 2 (Figure S1C)** | |
| **Intercept ($\beta_0$)** | $\beta = 0.14 \pm 0.07$<br>$t_{31} = 2.04$<br>$P = 0.050$ |
| **Performance/Evidence High confidence ($\beta_1$)** | $\beta = 12.23 \pm 1.49$<br>$t_{31} = 8.21$<br>$P = 2.810\times10^{-9}$ |
| **Performance/Evidence Low confidence ($\beta_2$)** | $\beta = 5.14 \pm 0.93$<br>$t_{31} = 5.50$<br>$P = 5.127\times10^{-6}$ |
| **Difference ($\beta_1 - \beta_2$)** | $t_{31} = 5.45$<br>$P = 5.854\times10^{-6}$ |

**Table S1 | Results of linear mixed-effects models for properties of confidence judgments**

#### Full behavioral GLMER

To assess whether our main behavioral results still hold in a full model, considering various other factors, we performed a model selection procedure of various linear mixed effect models. We used linear mixed-effects models (LMER) – as implemented in the lmer function from the lme4 package in R (Version 1.1-12; Bates, Maechler) (Bates et al., 2015).

We iteratively built several LMERs **(Table S2)**, and the final one was selected by model comparison, assessing model fit by using chi-square tests on the log-likelihood values, as well as comparison of the AIC and BIC model values **(Table S2)**. Model predictors were added whenever model fit was significantly improved.

The final model included fixed effects of incentive value (gain (1), neutral (0) or loss (-1)), evidence, accuracy (correct (1) or incorrect (0)), the interaction of accuracy end evidence, reaction time, and difficulty level (easy (1), medium (2), difficult (3)), as well as a random intercept and slope for the effect of incentive on confidence (model 9, see **Table S2**). Satterthwaite approximations (Schaalje et al., 2002) were used to calculate degrees of freedom and p-value estimates for the fixed effects' regression coefficients by using the 'lmerTest' package (Version 2.0-36, (Kuznetsova et al., 2017)). Visual inspection of residual plots did not reveal any obvious deviations from homoscedasticity or normality.

Final model results revealed that the significant effect of net incentive value on confidence still holds, while considering all other factors **(Table S3)**. Moreover, we found a significant effect of RT on confidence, showing that quicker choices lead to higher confidence levels. We also replicated that the link between confidence ratings and evidence is positive for correct and negative for incorrect responses.

| **Model** | **Model notation** | **AIC** | **BIC** | **Model comp.** | $\chi^2$ | **P-value** | **Winning model** |
|---|---|---|---|---|---|---|---|
| **1** | Confidence ~ Incentive + (1\|Subject) | 35083 | 34109 | | | | |
| **2** | Confidence ~ Incentive + (1+Incentive\|Subject) | 34077 | 34115 | 1 vs. 2 | 10.64 | 0.005 | 2 |
| **3** | Confidence ~ Incentive + Accuracy + (1+Incentive\|Subject) | 33920 | 33964 | 2 vs. 3 | 158.78 | $< 2\times10^{-16}$ | 3 |
| **4** | Confidence ~ Incentive + Accuracy + Evidence + (1+Incentive\|Subject) | 33817 | 33868 | 3 vs. 4 | 104.68 | $< 2\times10^{-16}$ | 4 |
| **5** | Confidence ~ Incentive + Accuracy*Evidence + (1+Incentive\|Subject) | 33767 | 33824 | 4 vs. 5 | 51.92 | $5.797\times10^{-13}$ | 5 |
| **6** | Confidence ~ Incentive + Accuracy*Evidence + Gender + (1+Incentive\|Subject) | 33769 | 33833 | 5 vs. 6 | 0.01 | 0.936 | 5 |
| **7** | Confidence ~ Incentive + Accuracy*Evidence + Age + (1+Incentive\|Subject) | 33769 | 33833 | 5 vs. 7 | 0.16 | 0.687 | 5 |
| **8** | Confidence ~ Incentive + Accuracy*Evidence + Difficulty + (1+Incentive\|Subject) | 33788 | 33808 | 5 vs. 8 | 33.11 | $6.458\times10^{-8}$ | 8 |
| 9 | Confidence ~ Incentive + RT + Accuracy*Evidence + Difficulty + (1+Incentive\|Subject) | 33142 | 33218 | 8 vs. 9 | 598.25 | $< 2\times10^{-16}$ | 9 |

**Table S2 | Model descriptions and comparison**
Shown here are the model notations of all nine models with their respective AIC and BIC values, as well as model comparisons with corresponding $\chi^2$ and P-values, resulting in the winning model 9.

| **Full behavioral GLMER Results**<br>***Confidence ~ Incentive + RT + Accuracy*Evidence + Difficulty + (1+Incentive\|Subject)*** | |
|---|---|
| **Intercept (B0)** | $\beta = 76.56 \pm 1.27$<br>$t_{45} = 60.36$<br>$P < 2\times10^{-16}$ |
| **Incentive** | $\beta = 0.88 \pm 0.30$<br>$t_{32} = 2.94$<br>$P = 0.006$ |
| **RT** | $\beta = -5.24 \pm 0.21$<br>$t_{4305} = -25.34$<br>$P < 2\times10^{-16}$ |
| **Accuracy** | $\beta = 3.30 \pm 0.42$<br>$t_{4290} = 7.86$<br>$P = 4.71\times10^{-15}$ |
| **Accuracy * Evidence** | $\beta = 2.83 \pm 0.50$<br>$t_{4275} = 5.69$<br>$P = 1.38\times10^{-8}$ |
| **Difficulty hard** | $\beta = -2.22 \pm 0.43$<br>$t_{4258} = -5.20$<br>$P = 2.07\times10^{-7}$ |
| **Difficulty medium** | $\beta = -1.53 \pm 0.41$<br>$t_{4256} = -3.71$<br>$P = 2.110\times10^{-4}$ |

**Table S3 | Results of general linear mixed-effects model**

Shown here are the results of the full linear mixed-effects model of the winning model. β: estimated regression coefficients for fixed effects ± estimated standard error of the regression coefficients, with corresponding t- and P-values.

#### Early certainty

In this section, we provide further details about the computation and properties of the early certainty variable. To verify that our model of early certainty is an appropriate proxy of confidence judgments, we performed similar behavioral analyses to confirm the three main properties of confidence judgments still hold for our early certainty variable. We performed identical analyses, substituting subjective confidence judgments for early certainty values.

Our results show that the measures of early certainty and performance are highly correlated ($R = 0.67 \pm 0.07$; **Figure S2A, Table S4**). Early certainty is also positively associated with evidence for correct trials, and negatively for incorrect trials (**Figure S2B, Table S4**). Finally, the relationship between performance and evidence is indeed higher in trials with high early certainty versus low early certainty (**Figure S2C, Table S4**).

When inspecting the beta values for the second model (**Figure S2C, Table S4**), we observed three statistical outliers (i.e. >1.5 times the interquartile range away from the 75$^{th}$ percentile) in the effect of evidence on performance in trials with high early certainty (**$\beta_1$**). These outliers were caused by the median-split of the early certainty trials into high and low variants, as these subjects performed (almost) perfectly in the high early certainty trials, causing the betas to inflate. Importantly, when excluding these subjects from the analyses, we found identical results, albeit stronger ($\beta_0 = 0.09 \pm 0.07$, $t_{28} = 1.26$, $P = 0.217$; $\beta_1 = 17.99 \pm 2.40$, $t_{28} = 7.49$, $P = 3.689\times10^{-8}$; $\beta_2 = 3.83 \pm 0.94$, $t_{28} = 4.06$, $P = 3.552\times10^{-4}$; Difference ($\beta_1 - \beta_2$): $t_{28} = 5.87$, $P = 2.595\times10^{-6}$).

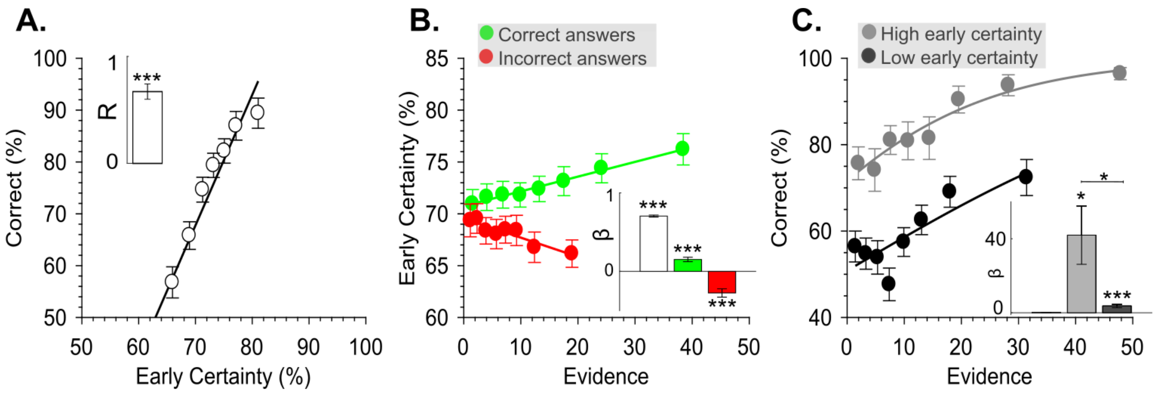

**Figure S2 | Properties of Early Certainty**

A: observed performance (% correct choices) as a function of early certainty. B: early certainty as function of evidence for correct (green) and incorrect (red) choices. C: observed performance (% correct choices) as a function of evidence, for high (gray) and low (black) early certainty trials. All plots include the three statistical outliers.

The insets presented on the side of each graph depict the results of the population-level analyses on the correlation coefficients (A) or on the regression coefficients (B and C). Error bars indicate inter-subject standard errors of the mean. *: P<0.05; **: P<0.01; ***P<0.001

| Model 1 (Figure S2B) | |
|---|---|
| **Intercept ($\beta_0$)** | $\beta = 0.70 \pm 0.01$<br>$t_{31} = 53.91$<br>$P = 3.304\times10^{-32}$ |
| **Confidence/Evidence Correct Answers ($\beta_1$)** | $\beta = 0.15 \pm 0.03$<br>$t_{31} = 5.45$<br>$P = 5.991\times10^{-6}$ |
| **Confidence/Evidence Incorrect Answers ($\beta_2$)** | $\beta = -0.27 \pm 0.05$<br>$t_{31} = -5.16$<br>$P = 1.370\times10^{-5}$ |
| **Model 2 (Figure S2C)** | |
| **Intercept ($\beta_0$)** | $\beta = 0.11 \pm 0.07$<br>$t_{31} = 1.51$<br>$P = 0.142$ |
| **Performance/Evidence High confidence ($\beta_1$)** | $\beta = 41.95 \pm 15.75$<br>$t_{31} = 2.67$<br>$P = 0.012$ |
| **Performance/Evidence Low confidence ($\beta_2$)** | $\beta = 3.64 \pm 0.90$<br>$t_{31} = 4.06$<br>$P = 3.052\times10^{-4}$ |
| **Difference ($\beta_1 - \beta_2$)** | $t_{31} = 2.40$<br>$P = 0.023$ |

**Table S4 | Results of linear mixed-effects models for properties of early certainty**

Moreover, to validate that our model of early certainty correlates highly with subjective confidence and choice and stimulus features, but does not show a statistical relationship with incentives, we built a linear mixed-effects model using the lme4 package in R. We used early certainty as dependent variable and added RT, accuracy, evidence and the interaction between evidence and accuracy as predictors. Indeed, the results showed that RT, accuracy and the accuracy * evidence interaction all significantly contributed to early certainty, while no effect of incentive value on early certainty was found (**Table S5)**.

| **Early Certainty GLMER Results**<br><br>***Early Certainty ~ Incentive + RT + Accuracy*Evidence + (1\|Subject)*** | |
|---|---|
| **Intercept (B0)** | β = 75.52 ± 1.15<br>$t_{33}$ = 65.63<br>$P < 2\times10^{-16}$ |
| **Incentive** | β = 0.07 ± 0.08<br>$t_{4288}$ = 0.84<br>P = 0.404 |
| **RT** | β = -5.69 ± 0.08<br>$t_{4292}$ = -71.90<br>$P < 2\times10^{-16}$ |
| **Accuracy** | β = 3.61 ± 0.16<br>$t_{4288}$ = 22.75<br>$P < 2\times10^{-16}$ |
| **Accuracy * Evidence** | β = 2.50 ± 0.19<br>$t_{4288}$ = 13.16<br>$P < 2\times10^{-16}$ |

**Table S5 | Results of general linear mixed-effects model**
Shown here are the results of the full linear mixed-effects model. β: estimated regression coefficients for fixed effects ± estimated standard error of the regression coefficients, with corresponding t- and P-values.

#### Activation Tables for GLM1 and GLM3 (Figure 3).

| GLM1 | | | | | | | | |
|---|---|---|---|---|---|---|---|---|
| **Effect** | **Brain Region** | **k** | **Peak z-score** | **P (cluster FWE corrected)** | **Peak voxel MNI coordinates** | | | |
| **Early certainty +** | VMPFC | 95 | 3.97 | 0.004 | 3 | 29 | -7 | LR |
| | | | | | -9 | 65 | -4 | LR |
| | | | | | -3 | 38 | -7 | LR |
| | PPC | 59 | 3.79 | 0.03 | -3 | -43 | 32 | LR |
| | | | | | 6 | -52 | 32 | LR |
| | | | | | -3 | -58 | 29 | LR |
| **Early certainty -** | Insula<br>Inferior frontal gyrus<br>RLPFC / DLPFC<br>Putamen | 873 | 5.92 | <0.001 | 33 | 20 | 8 | R |
| | | | | | 45 | 14 | 2 | R |
| | | | | | 45 | 38 | -4 | R |
| | DLPFC<br>RLPFC | 176 | 4.55 | <0.001 | -42 | 26 | 35 | L |
| | | | | | -30 | 47 | 11 | L |
| | | | | | -36 | 29 | 26 | L |
| | Supplementary motor area<br>Mid cingulate cortex<br>Anterior cingulate cortex | 583 | 5.51 | <0.001 | 3 | 11 | 47 | LR |
| | | | | | 6 | 23 | 35 | LR |
| | | | | | 9 | 17 | 41 | LR |
| | Supramarginal gyrus | 76 | 4.45 | 0.011 | 51 | -19 | 20 | R |
| | | | | | 42 | -22 | 23 | R |
| | Posterior Insula | | | | 36 | -13 | 20 | R |
| | Inferior Occipital Gyrus | 57 | 4.33 | 0.034 | -48 | -70 | -1 | L |
| | | | | | -42 | -61 | -7 | L |
| | Inferior parietal lobe | 332 | 4.99 | <0.001 | -48 | -34 | 41 | L |
| | | | | | -30 | -52 | 41 | L |
| | Postcentral gyrus | | | | -57 | -22 | 35 | L |
| | Anterior insula | 170 | 4.95 | <0.001 | -33 | 17 | 5 | L |
| | | | | | -33 | 26 | -7 | L |
| | Cerebellum | 121 | 4.98 | 0.001 | -36 | -55 | -31 | L |
| | | | | | -15 | -43 | -25 | L |
| **Incentive +** | VMPFC | 46 | 4.32 | 0.011 | 3 | 47 | -4 | LR |
| | Anterior medial prefrontal cortex / DMPFC | 36 | 4.14 | 0.032 | 0 | 56 | 11 | LR |
| | DLPFC | 39 | 4.01 | 0.023 | -27 | 38 | 55 | L |
| | | | | | -6 | 35 | 32 | L |
| **Incentive -** | Angular gyrus | 54 | 4.06 | 0.005 | 39 | -55 | 41 | R |
| | | | | | 39 | -58 | 50 | R |
| | Superior occipital lobe | | | | 27 | -61 | 38 | R |
| **Confidence +** | Cerebellum<br>Lingual gyrus (visual cortex) | 997 | 6.19 | <0.001 | 12 | -52 | -16 | R |
| | | | | | 18 | -70 | -13 | R |
| | | | | | 21 | -70 | -4 | R |
| | Putamen | 328 | 4.83 | <0.001 | -33 | -10 | -1 | L |
| | | | | | -30 | 2 | 5 | L |

| | | | | | | | | |
|---|---|---|---|---|---|---|---|---|
| | | | | | -42 | -8 | 11 | L |
| | Primary motor cortex | 244 | 4.88 | <0.001 | -33 | -28 | 59 | L |
| | | | | | -42 | -22 | 41 | L |
| | | | | | -42 | -19 | 56 | L |
| | Anterior cingulate cortex | 90 | 4.55 | 0.001 | -6 | 23 | 29 | L |
| | | | | | 3 | 20 | 26 | R |
| | Mid cingulate cortex | | | | -6 | 2 | 38 | LR |
| | Parahippocampal gyrus | 64 | 4.01 | 0.008 | -24 | -37 | -13 | L |
| | Fusiform gyrus | | | | -24 | -46 | -10 | L |
| | Middle temporal gyrus | 56 | 3.97 | 0.016 | 48 | -70 | 2 | R |
| | | | | | 48 | -52 | 17 | R |
| | | | | | 51 | -64 | 17 | R |
| | Precuneus | 75 | 4.29 | 0.004 | -15 | -49 | 8 | L |
| | | | | | -6 | -58 | 23 | L |
| | | | | | -9 | -58 | 14 | L |
| **Confidence -** | Lingual gyrus (visual cortex) | 302 | 5.82 | <0.001 | -15 | -82 | -4 | L |
| | Cerebellum | | | | -15 | -55 | -16 | L |
| | Primary motor cortex | 55 | 4.26 | 0.018 | 39 | -19 | 59 | R |
| | | | | | 36 | -19 | 44 | R |
| | | | | | 57 | -16 | 44 | R |

**Table S6 | GLM1 activation table**

Brain activations (whole brain analyses) of GLM1 showing activity related to early certainty at choice moment, as well as activity related to incentive and confidence at incentive/rating moment. All whole-brain activation maps were thresholded using family-wise error correction for multiple correction (FWE) at cluster level ($P_{FWE\_clu} < 0.05$), with a voxel cluster-defining threshold of $P<0.001$ uncorrected. Activity that positively correlates to given variable is denoted by '+', whereas negative correlations are denoted by '-'.

| **GLM 3** | | | | | | | | |
|---|---|---|---|---|---|---|---|---|
| **Effect** | **Brain Region** | **k** | **Peak z-score** | **P (cluster FWE corrected)** | **Peak voxel MNI coordinates** | | | |
| **Expected Value +** | VMPFC<br>Anterior medial prefrontal cortex / DMPFC<br>Anterior cingulate cortex (dorsal + ventral) | 336 | 4.93 | <0.001 | 0 | 47 | -4 | LR |
| | | | | | 0 | 56 | 11 | LR |
| | | | | | 0 | 32 | 11 | LR |
| | Insula | 37 | 4 | 0.038 | -36 | 5 | 1 | L |
| | | | | | -36 | -1 | 5 | L |

**Table S7 | GLM3 activation table**

Brain activations (whole brain analyses) of GLM3 showing activity related to EV at incentive/rating moment. All whole-brain activation maps were thresholded using family-wise error correction for multiple correction (FWE) at cluster level ($P_{FWE\_clu} < 0.05$), with a voxel cluster-defining threshold of $P<0.001$ uncorrected. Activity that positively correlates to given variable is denoted by '+'.

### Explorative analyses dACC

***Explorative analysis of dACC results show overlap between confidence and EV signal***

While we did not find clear evidence for VMPFC activity correlating with confidence at our pre-specified statistical threshold, we did find a cluster of dACC activity positively correlating with both confidence (**Figure 2A**) and EV (**Figure 2B**). We therefore applied our ROI analytical strategy – originally designed for the VMPFC – to the dACC. Like for the VMPFC analyses we built an independent anatomical ROI of the dACC from the Brainnetome Atlas (Fan et al., 2016) (**Figure S3A**).

We compared early certainty, incentive and confidence-related activations during both time-points in all available GLMs within the dACC ROI (see **Figure 4** in main text for comparable analysis in VMPFC). Thus, we extracted individual standardized regression coefficients (t-values) from the dACC, corresponding to these respective activations and statistically compared them using repeated measure ANOVAs and post-hoc paired t-tests (**Figure S3, Table S8**). Activations for early certainty during choice moment were similar for all GLMs (ANOVA $F(4,29) = 1.75$, $P = 0.144$; **Figure S3B**), and all were significantly negatively related to early certainty (GLM1: $t_{29} = -2.48$, $P = 0.019$; GLM2a: $t_{29} = -2.48$, $P = 0.019$; GLM2b: $t_{29} = -2.39$, $P = 0.024$; GLM3: $t_{29} = -2.48$, $P = 0.019$; GLM4: $t_{29} = -2.51$, $P = 0.018$). GLM specification had an impact on the incentive activation (ANOVA, main effect of GLM; $F(3,29) = 19.13$, $P = 1.292\times10^{-9}$; **Figure S3C**), but not on the confidence activations (ANOVA, main effect of GLM; $F(3,29) = 1.95$, $P = 0.127$; **Figure S3D**) during incentive/rating moment. In the incentive case, post-hoc t-tests showed that T-values extracted from the GLM3 that related to the EV regressor were significantly higher than from other GLMs with a different coding of incentives (GLM1 versus GLM3: $t_{29} = -5.22$, $P = 1.378\times10^{-5}$; GLM2b versus GLM3: $t_{29} = -4.45$, $P = 1.164\times10^{-4}$; GLM4 versus GLM3: $t_{29} = -4.31$, $P = 1.708\times10^{-4}$), but activity related to EV and confidence or certainty during rating moment were found to be similarly strong.

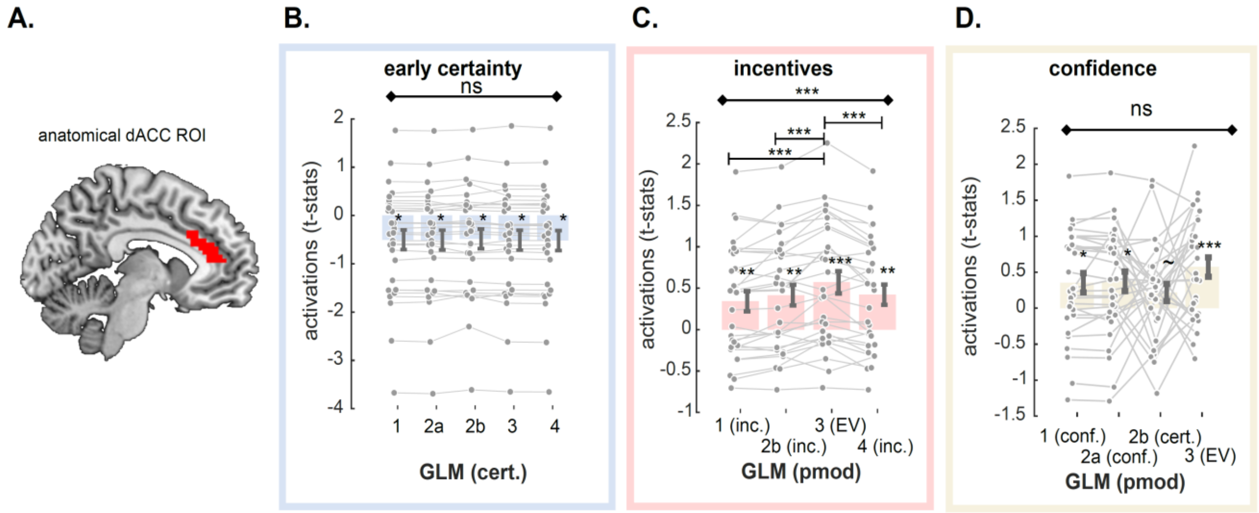

**Figure S3 | A.** Anatomical dACC region of interest (ROI). **B-D** Comparison of dACC activations to different specifications of early certainty during choice moment (B), incentives during incentive/rating moment (C) and confidence during incentive/rating moment (D), as implemented in the different GLMs. Dots represent individual activations; bar and error bars indicate sample mean ± sem. Grey lines highlight within subject variation across the different specifications.
Cert: early certainty; Inc.: incentives; conf: confidence; EV: expected value;
Diamond-ended horizontal bars indicate the results of repeated-measure ANOVAs. Dash-ended horizontal bars indicate the result of post-hoc paired t-tests.
~ P < 0.10; * P<0.05; ** P<0.01; *** P < 0.001

Finally, we repeated the qualitative falsification exercise (see **Figure 5** in the main text) for the dACC ROI. We extracted the dACC activations for all regressors in GLM5 using our ROI, and compared them with the theorized qualitative patterns (**Figure S4, Table S9-10**). At the stimulus/choice moment, we found no effect of incentive conditions on dACC baseline activity, nor on its correlation with confidence – "slope" (ANOVA baseline: $F(2,29) = 0.05$, $P = 0.952$; ANOVA slope: $F(2,29) = 0.63$, $P = 0.534$). At rating moment, incentive conditions had an effect on dACC baseline activity (ANOVA $F(2,29) = 12.30$, $P = 3.527\times10^{-5}$). Post-hoc testing revealed that dACC baseline activity was significantly positive in all incentive conditions (Loss: $t_{29} = 3.96$, $P = 4.469\times10^{-4}$; Neutral: $t_{29} = 2.69$, $P = 0.012$; Gain: $t_{29} = 6.31$, $P = 6.807\times10^{-7}$), but larger in gain versus loss ($t_{29} = -3.63$, $P = 0.001$) and in gain vs neutral conditions ($t_{29} = -4.10$, $P = 3.009\times10^{-4}$), but not in loss vs neutral condition ($t_{29} = 1.71$, $P = 0.098$) (see **Table S9-10**). Incentive conditions had a marginally significant effect (ANOVA $F(2,29) = 3.12$ $P = 0.052$) on the slope of the correlation of dACC activity with confidence, where only in the gain condition the slope was positive ($t_{29}$ = 3.35, P = 0.002). Post-hoc testing showed that the correlation with confidence was only significantly higher in gain versus loss ($t_{29}$ = -2.37, P = 0.025), and marginally higher for gain versus neutral conditions ($t_{29}$ = -1.95, P = 0.060), whereas no difference was found for neutral versus loss condition ($t_{29}$ = -0.18, P = 0.860). Again, similar to the results in the VMPFC, the observed pattern of dACC activity was not featured in the EV model, nor in the confidence model, or any other model prediction, and thus points to a more complex picture of disruption of metacognitive signals due to motivational signals.

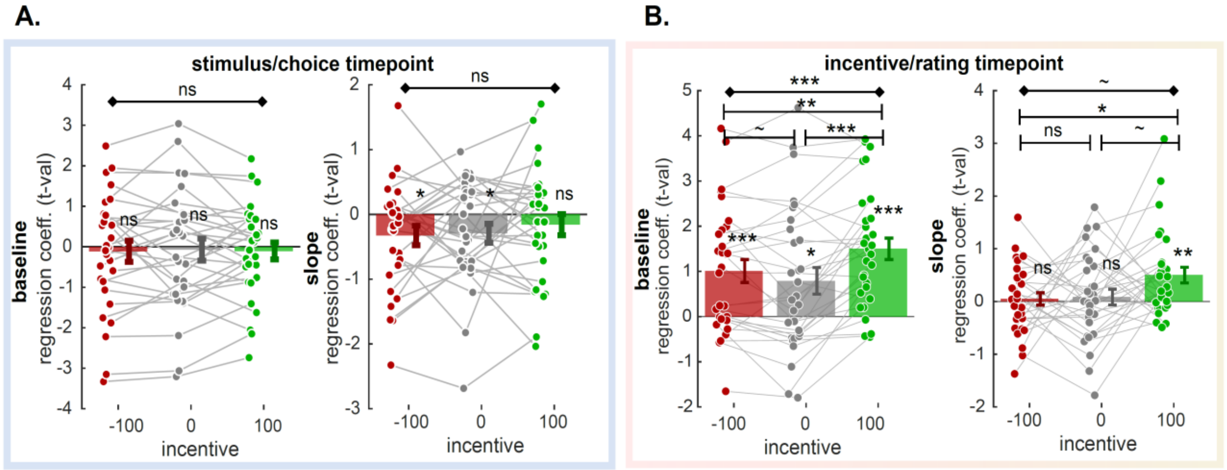

**Figure S4 | A-B.** dACC ROI analysis. T-values corresponding to baseline and regression slope were extracted in the three incentive conditions, and at the two time-points of interest (A: stimulus/choice; B: incentive/rating). Dots represent individual activations; bar and error bars indicate sample mean ± sem. Grey lines highlight within subject variation across the different incentive conditions.
Diamond-ended horizontal bars indicate the results of repeated-measure ANOVAs. Dash-ended horizontal bars indicate the result of post-hoc paired t-tests.
~ P < 0.10; * P<0.05; ** P<0.01; *** P < 0.001

| | | | | | |
|---|---|---|---|---|---|
| **Early certainty** | **GLM1** | **GLM2a** | **GLM2b** | **GLM3** | **GLM4** |
| | -0.50 ± 0.2<br>$t_{29}$ = -2.48<br>P = 0.019 | -0.51 ± 0.2<br>$t_{29}$ = -2.48<br>P = 0.019 | -0.48 ± 0.2<br>$t_{29}$ = -2.39<br>P = 0.024 | -0.51 ± 0.21<br>$t_{29}$ = -2.48<br>P = 0.019 | -0.52 ± 0.21<br>$t_{29}$ = -2.51<br>P = 0.018 |
| | **ANOVA (Main effect of GLM)** | - | - | - | - |
| | F(4,29) = 1.75<br>P = 0.144 | - | - | - | |
| **Incentive** | | **GLM1** | **GLM2b** | **GLM3** | **GLM4** |
| | | 0.34 ± 0.12<br>$t_{29}$ = 2. 82<br>P = 0.009 | 0.41 ± 0.12<br>$t_{29}$ = 3. 34<br>P = 0.002 | 0.57 ± 0.13<br>$t_{29}$ = 4. 25<br>P = $2.022\times10^{-4}$ | 0.42 ± 0.12<br>$t_{29}$ = 3. 43<br>P = 0.002 |
| | **ANOVA (Main effect of GLM)** | **T-Test (3 vs 1)** | **T-Test (3 vs 2b)** | - | **T-Test (3 vs 4)** |
| | F(3,29) = 19.13<br>P= $1.292\times10^{-9}$ | -0.23 ± 0.09<br>$t_{29}$ = -5.22<br>P = $1.378\times10^{-5}$ | -0.16 ± 0.07<br>$t_{29}$ = -4.45<br>P = $1.164\times10^{-4}$ | - | -0.15 ± 0.07<br>$t_{29}$ = -4.31<br>P = $1.708\times10^{-4}$ |
| **Confidence** | | **GLM1** | **GLM2a** | **GLM2b** | **GLM3** |
| | | 0. 35 ± 0.13<br>$t_{29}$ = 2. 65<br>P = 0. 013 | 0.37 ± 0.14<br>$t_{29}$ = 2. 75<br>P = 0. 010 | 0.22 ± 0.12<br>$t_{29}$ = 1.82<br>P = 0. 080 | 0.57 ± 0.13<br>$t_{29}$ = 4. 25<br>P = $2.022\times10^{-4}$ |
| | **ANOVA (Main effect of GLM)** | **T-Test (3 vs 1)** | **T-Test (3 vs 2a)** | **T-Test (3 vs 2b)** | - |
| | F(3,29) = 1.95<br>P = 0.127 | -0.22 ± 0.36<br>$t_{29}$ = -1. 24<br>P = 0. 226 | -0.20 ± 0.35<br>$t_{29}$ = -1.15<br>P = 0.258 | -0.35 ± 0.33<br>$t_{29}$ = -2.20<br>P = 0.036 | |

**Table S8 | Comparison of ACC parametric activity (t-values) as a function of model specification (GLMs)**

The table reports descriptive and inferential statistics on ACC ROI parametric activations with three different variables of interest: early certainty effects at choice moment, incentive effects at rating moment and confidence effects at rating moment (see **Figure 4**). Per effect of interest, results of one-sample t-tests against zero, repeated-measure (RM) ANOVAs on the main effect of GLMS, and post-hoc t-test results are shown.

| Choice/Stim | | | | | |
|---|---|---|---|---|---|
| | **baseline** | **Inc. -100** | **Inc. 0** | **Inc. +100** | **ANOVA** |
| | | -0.11 ± 0.26<br>$t_{29}$ = -0.43<br>P = 0.670 | -0.07 ± 0.27<br>$t_{29}$ = -0.26<br>P = 0.796 | -0.10 ± 0.21<br>$t_{29}$ = -0.49<br>P = 0.626 | F(2,29) = 0.05<br>P = 0.952 |
| | **slope** | **Inc. -100** | **Inc 0** | **Inc. +100** | **ANOVA** |
| | | -0.33 ± 0.15<br>$t_{29}$ = -2.17<br>P = 0.038 | -0.29 ± 0.15<br>$t_{29}$ = -1.98<br>P = 0.057 | -0.16 ± 0.16<br>$t_{29}$ = -0.98<br>P = 0.337 | F(2,29) = 0.63<br>P = 0.534 |

**Table S9 | Comparison of ACC activity at choice moment (t-values), as a function of incentive condition**

The table reports descriptive and inferential statistics on ACC ROI parametric activations in our three incentive conditions during choice moment, for both baseline activity as well as the correlation with early certainty (i.e. slope) (see **Figure S4**). Results of RM ANOVAs and one-sample t-tests against 0 are shown.

| Incentive/rating | | Inc -100 | Inc 0 | Inc +100 | ANOVA |
|---|---|---|---|---|---|
| | baseline | 1.01 ± 0.25<br>$t_{29}$ = 3.96<br>P = 4.469×$10^{-4}$ | 0.79 ± 0.29<br>$t_{29}$ = 2.69<br>P = 0.012 | 1.50 ± 0.24<br>$t_{29}$ = 6.31<br>P = 6.807×$10^{-7}$ | F(2,29) = 12.30<br>P = 3.527×$10^{-5}$ |
| | | **T-Test**<br>[-100 vs 0] | **T-Test**<br>[0 vs 100] | **T-Test**<br>[-100 vs 100] | |
| | | 0.22 ± 0.13<br>$t_{29}$ = 1.71<br>P = 0.098 | -0.71 ± 0.17<br>$t_{29}$ = -4.10<br>P = 3.009×$10^{-4}$ | -0.49 ± 0.14<br>$t_{29}$ = -3.63<br>P = 0.001 | |
| | | **Inc -100** | **Inc 0** | **Inc +100** | **ANOVA** |
| | slope | 0.05 ± 0.12<br>$t_{29}$ = 0.41<br>P = 0.685 | 0.08 ± 0.15<br>$t_{29}$ = 0.55<br>P = 0.583 | 0.50 ± 0.15<br>$t_{29}$ = 3.35<br>P = 0.002 | F(2,29) = 3.12<br>P = 0.052 |
| | | **T-Test**<br>[-100 vs 0] | **T-Test**<br>[0 vs 100] | **T-Test**<br>[-100 vs 100] | |
| | | -0.04 ± 0.20<br>$t_{29}$ = -0.18<br>P = 0.860 | -0.42 ± 0.21<br>$t_{29}$ = -1.95<br>P = 0.060 | -0.45 ± 0.19<br>$t_{29}$ = -2.37<br>P = 0.025 | |

**Table S10 | Comparison of ACC activity at rating moment (t-values), as a function of incentive condition**

The table reports descriptive and inferential statistics on ACC ROI parametric activations in our three incentive conditions during rating moment, for both baseline activity as well as the correlation with confidence (i.e. slope) (see **Figure 5C**). Results of one-sample t-tests against 0, RM ANOVAs and post-hoc t-tests are shown.

#### References

Bates, D., Mächler, M., Bolker, B., & Walker, S. (2015). Fitting Linear Mixed-Effects Models Using lme4. *Journal of Statistical Software*, *67*(1), 1–48. https://doi.org/10.18637/jss.v067.i01

Fan, L., Li, H., Zhuo, J., Zhang, Y., Wang, J., Chen, L., Yang, Z., Chu, C., Xie, S., Laird, A. R., Fox, P. T., Eickhoff, S. B., Yu, C., & Jiang, T. (2016). The Human Brainnetome Atlas: A New Brain Atlas Based on Connectional Architecture. *Cerebral Cortex*, *26*(8), 3508–3526. https://doi.org/10.1093/cercor/bhw157

Kuznetsova, A., Brockhoff, P. B., & Christensen, R. H. B. (2017). lmerTest Package: Tests in Linear Mixed Effects Models. *Journal of Statistical Software*, *82*(13), 1–26. https://doi.org/10.18637/jss.v082.i13

Lebreton, M., Langdon, S., Slieker, M. J., Nooitgedacht, J. S., Goudriaan, A. E., Denys, D., Holst, R. J. van, & Luigjes, J. (2018). Two sides of the same coin: Monetary incentives concurrently improve and bias confidence judgments. *Science Advances*, *4*(5), eaaq0668. https://doi.org/10.1126/sciadv.aaq0668

Sanders, J. I., Hangya, B., & Kepecs, A. (2016). Signatures of a Statistical Computation in the Human Sense of Confidence. *Neuron*, *90*(3), 499–506. https://doi.org/10.1016/j.neuron.2016.03.025

Schaalje, G. B., McBride, J. B., & Fellingham, G. W. (2002). Adequacy of approximations to distributions of test statistics in complex mixed linear models. *Journal of Agricultural, Biological, and Environmental Statistics*, *7*(4), 512. https://doi.org/10.1198/108571102726
